# Activation of ERK by altered RNA splicing in cancer

**DOI:** 10.1101/2022.08.31.505957

**Authors:** Yushan Zhang, Md Afjalus Siraj, Prabir Chakraborty, Robert Tseng, Li-Ting Ku, Shamik Das, Anindya Dey, Shailendra Kumar Dhar Dwivedi, Geeta Rao, Min Zhang, Da Yang, Md Nazir Hossen, Wei-Qun Ding, Kar-Ming Fung, Resham Bhattacharya, Luisa Escobar-Hoyos, Priyabrata Mukherjee

## Abstract

Many cancers carry change-of-function mutations affecting RNA splicing factors, however, less is known about the functional consequences of upregulated RNA splicing factors in cancer. Here, we demonstrate that SMNDC1, a poorly studied splicing factor, which we found to be upregulated in multiple carcinomas and associated with poor patient prognosis, promotes cell proliferation, clonal expansion, and tumor growth by promoting the retention of G-rich exons, which otherwise would be excluded or retained at a lower rate after RNA splicing in normal cells. Inclusion of exon 4 (E4) of *MAPK3* (ERK1), which encodes both kinase phosphorylation sites (Thr202/Tyr204), was among the promoted exons by SMNDC1. Forced exclusion of *MAPK3-*E4 using anti-sense oligos inhibited the ERK1 phosphorylation, expression of target genes and decreased tumor cell growth. These data support that cancer cells exploit a “splicing switch” to promote ERK kinase activity and offer a druggable alternative to block oncogenic signaling and altered RNA splicing in cancer cells

**SIGNIFICANCE:** ERK signaling promotes tumor growth and survival. Exon 4 of *MAPK3* (ERK1) encodes the activation phosphorylation sites of ERK1 kinase. Aberrant RNA splicing induced by SMNDC1 in cancer cells increases the retention of exon 4 during mRNA splicing, unleashes the kinase activity. SMNDC1 potentializes as a cancer therapeutic target.

## INTRODUCTION

RNA splicing is a co-transcriptional process that removes and joins exons and introns from precursor messenger RNA (pre-mRNA) to generate the mature mRNA (1, 2). This process occurs in nucleus and is catalyzed by the spliceosome, a dynamic megadalton complex comprising five small nuclear ribonucleoprotein particles (snRNPs) U1, U2, U4, U5 and U6, and more than 300 splicing factors and regulators (3). Recurrent mutations affecting RNA splicing factors commonly occur in many hematologic malignancies (4). In solid malignancies, however, it is more common to find altered expression of RNA splicing factors and regulators, compared to normal/benign counterparts (4). Despite these findings, the roles of these up- or down-regulated splicing proteins in carcinomas is poorly understood. Survival motor neuron domain containing 1 (SMNDC1), also known as splicing factor 30 (SPF30), was identified as a nuclear protein and an splicing regulator by mass spectrometry of cervical cancer cell lines and expressed-sequence tagged-database searching (5). It was further demonstrated that SMNDC1 associates with both U4/U5/U6 and U2 snRNP components (6-8), is differentially expressed among human normal tissues (9) and upregulated at the transcriptional level in cancers of the colon, breast, kidney, lung, uterus, and blood (10, 11).

Despite of the unknown role of altered expression of RNA splicing factors, on the other hand, it is well known that the dysfunction of the EGFR-RAS-RAF-ERK pathway is a major trigger for the development of most cancer types (12). Activation of downstream ERK is mainly triggered by activating mutations in upstream proteins such as EGFR, RAS, and RAF (found in 50% of tumors) (13) and by altered RNA splicing of coding exons in these same genes (14-20), leading to deregulated proliferation and differentiation in cancer cells. ERK1 and ERK2 are the two most important downstream members of the EGFR-RAS-RAF-ERK pathway, with molecular weights of 44 and 42 kDa, respectively (21). ERK1/2 are serine/threonine protein kinases that are generally located in the cytoplasm and upon activation by phosphorylation in key residues, they enter the nucleus and regulate transcription factor activity and expression of many genes including cyclin D and ELK1 (21). Here, we report a novel mechanism by which cancer cells activate ERK1 signaling and is dependent on the overexpression of splicing regulator SMNDC1, which increases the retention of cassette exon that encodes both kinase phosphorylation sites (Thr202/Tyr204). These data support a model by which cancer cells exploit a “splicing switch” to promote ERK1 activity and offer a druggable alternative to block oncogenic signaling and altered RNA splicing in cancer cells.

## RESULTS

### SMNDC1 expression is upregulated in pancreatic and ovarian cancers and associated with negative patient outcome

We previously identified SMNDC1 as a cancer-specific protein using a nanoparticle proteomic screen comparing normal versus malignant ovarian cells (22). Following this approach, we identified that SMNDC1 is also enriched in pancreatic ductal adenocarcinoma (PDAC) cells (**Supplementary Fig. S1A**). To further investigate the association of SMNDC1 expression with cancer, we performed genomic and proteomic *in silico* analyses using Oncomine (23) and found that expression of SMNDC1 is significantly upregulated in both PDACs and ovarian carcinomas (OvCa) when compared to the corresponding normal/benign tissues in six patient cohorts with RNA expression (**Fig. 1A**). Furthermore, using “The Cancer Genome Atlas” we found that 16% of PDACs and 10% OvCas had higher mRNA expression of SMNDC1, and this overexpression is associated with worst patient survival in PDAC, OvCa, and other carcinomas (**Fig. 1B-C, Supplementary Fig. S1B-C**). To corroborate these findings and further understand the specificity of SMNDC1 expression in cancer cells within clinical specimens, we performed immunohistochemistry using a tissue microarray (TMA, **Supplementary Table ST1**) that contained 90 PDACs and 71 benign adjacent pancreatic tissues. After pathology scoring, it was found that SMNDC1 expression is doubled in the nuclei of PDAC cells versus the benign pancreatic cells and its expression is associated to increased tumor grade and poor patient outcome (**Fig. 1D-G, Supplementary Fig. S1D-F**). These results suggest that SMNDC1 expression is associated with malignant transformation of both pancreatic and ovarian cells, and it is a negative prognostic biomarker.

**Figure 1.**
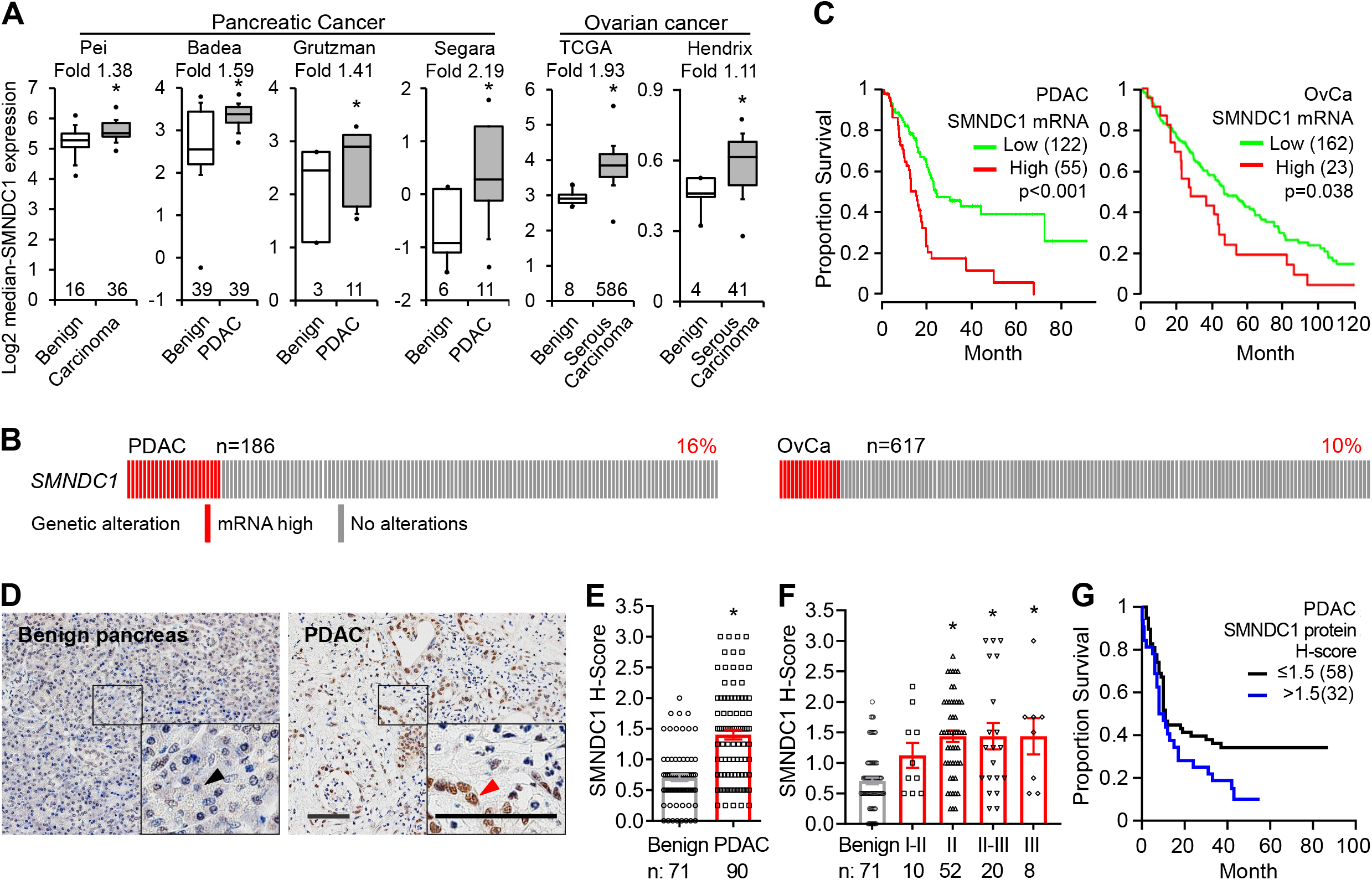
SMNDC1 expression in and association with cancer. **A**, Oncomine mRNA expression datasets of pancreatic ductal adenocarcinoma (PDAC) and Ovarian cancer (OvCa). The box and whisker plots showed the minimum, 25% quartile, median, 75% quartile, the maximum and outliers. Number at the bottom: number of cases. **B**, Oncoprint visualizing SMNDC1 genetic alteration in PDAC and OvCa. n: case number. **C**, Association of SMNDC1 mRNA expression with cancer survival. PDAC curves are based on TCGA database; OvCa curves are based on Gene Expression Omnibus (GEO) dataset (GSE26712). **D**, Example IHC images showing SMNDC1 protein expression in PDAC tissue and benign pancreatic tissue. Arrow heads: cancer cell nucleus (red) or benign epithelial cell nucleus (black). Scale bar: 100 µm. **E**, TMA H-score for PDAC. n, case number. **F**, Association of SMNDC1 protein expression with tumor grade based on PDAC TMA staining. **G**, Association of SMNDC1 protein expression with the survival of PDAC patients based on TMA H-score. H-score cutoff = 1.5. *, p<0.05.

### SMNDC1 promotes cell proliferation, colony formation and enhances *in vivo* tumor growth

To determine the function of SMNDC1 at the cellular level in cancer, we first investigated whether depletion of SMNDC1 inhibits cancer cell growth *in vitro*. Western blotting showed that SMNDC1 protein expression is upregulated in PDAC and OvCa cell lines compared to their normal counterparts─ human pancreatic ductal epithelial cell line (HPDEC) or ovarian surface epithelial cell line (OSE) (**Fig. 2A**), reinforcing the results from clinical samples that show upregulation of SMNDC1 in malignancy. SMNDC1 expression in three PDAC cell lines and two OvCa cell lines was transiently or stably knocked down with RNAi (**Fig. 2B, Supplementary Fig. S2A**), and cell proliferation and colony formation were examined. Knockdown of SMNDC1 by either siRNA or shRNAs significantly inhibited proliferation (**Fig. 2C, Supplementary Fig. S2B-C**) and colony formation in all cell lines tested (**Fig. 2D-E, Supplementary Fig. S2D-G**). Likewise, overexpression of SMNDC1 increased cell proliferation (**Fig. 2F-G**), suggesting the on-target effect of this protein in cancer. To extend our observations to mouse xenograft models, we subcutaneously inoculated PDAC and OvCa cell lines with deficient and proficient SMNDC1 expression into mice and monitored tumor growth. Size and weight of tumors deficient for SMNDC1 were significantly smaller than in the control groups (**Fig. 2H-I, Supplementary Fig. S2H-I**). To determine the cellular mechanism that led to decreased tumor growth upon knockdown of SMNDC1, we performed stains in tumors for markers of proliferation (Ki67) and cell death (tunnel stain) and found that tumors proficient for SMNDC1 had increased cell cycle activity and decreased cell death compared to SMNDC1-deficient group (shSMNDC1) (**Fig. 2J-K, Supplementary Fig. S2J**). Taken together, both the *in vitro* and *in vivo* results evidenced the role of SMNDC1 in promoting increased cell proliferation and tumor growth.

**Figure 2.**
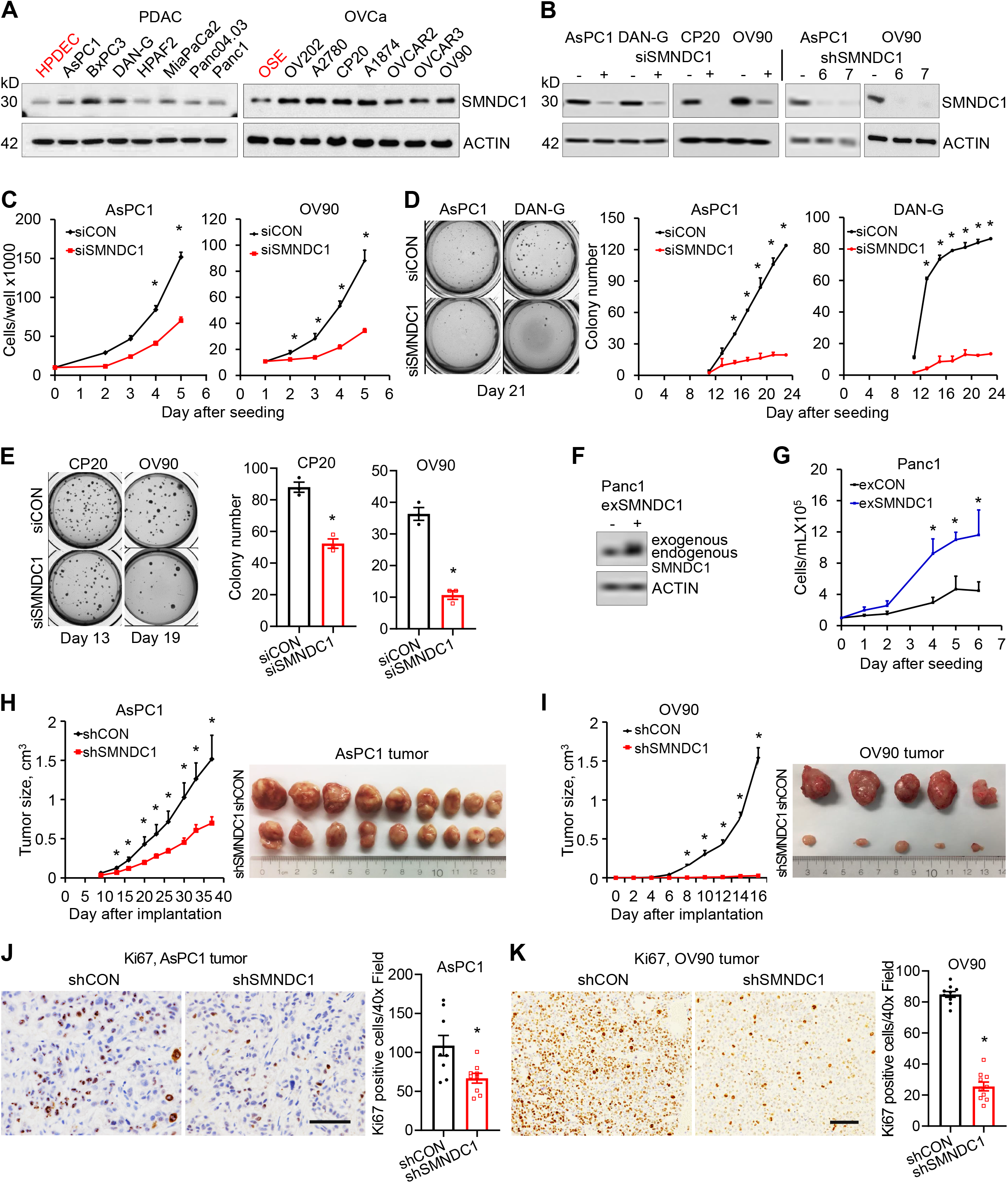
SMNDC1 promotes cancer cell proliferation, colony formation and *in vivo* tumor growth in mice. **A**. SMNDC1 protein expression in PDAC and OvCa cell lines and their normal counterparts HPDEC and OSE. ACTIN was used as a loading control. **B**, Efficiency of SMNDC1 knockdown by siRNA and shRNA in PDAC and OvCa HGSOC cell line (OV90) and UC cell line (CP20). shRNAs targeting two different sequences of SMNDC1, designated as shSMNDC1-6 and shSMNDC1-7, were used. **C**, Proliferation of cancer cells upon transient silencing of SMNDC1. **D, E**, 3D colony formation (colony number) of PDAC cells (**D**) or OvCa cells (**E**) upon transient silencing of SMNDC1. **F**, SMNDC1 was stably overexpressed in Panc1 cells (exSMNDC1). **G**, Proliferation of cancer cells with stable overexpression of SMNDC1. **H, I**, SMNDC1 stable knockdown in AsPC1 cells (**H**) or OV90 cells (**I**) were inoculated subcutaneously into the flank of nude mice (n=9 for AsPC1, n=10 for OV90). Tumor sizes were measured twice a week (AsPC1) or every other day (OV90). Tumor images show the size of tumors at experiment termination. **J, K**, Xenograft tumor tissue Ki67 staining and quantification. Scale bar: 100 µm. *, p<0.05.

### SMNDC1 upregulates the ERK pathway and blocks pro-apoptotic pathways

To gain mechanistic insight on the function of SMNDC1, we screened the phosphorylation state of multiple key cell signaling molecules linked to cell proliferation, survival and apoptosis (24, 25), and examined whether silencing SMNDC1 alters the activity of related molecules. Phosphorylation of ERK1/2 (T202/Y204, T185/Y187), AKT1/2/3 (S473), CREB (S133), CHK2 (T68), CJUN (S63) and WNK1 (T60) was found to be significantly inhibited in cells with SMNDC1 knockdown (**Fig. 3A**). Given that phosphorylation changes of ERK1/2, AKT1/2/3 and CREB are critical in cell apoptosis, survival and proliferation pathways (26, 27), we confirmed these changes by western blotting (**Supplementary Fig. S3A-B**). Among these kinases, ERK1 phosphorylation and downstream transcriptional target genes (Cyclin D and ELK1) were the most affected by SMNDC1 loss (**Fig. 3B-E**). These results suggested that SMNDC1 regulates apoptosis, survival and proliferation pathways, through kinases such as ERK1/2, AKT1/2/3 and CREB. To determine if these phosphorylation changes led to the activation or deactivation of proliferation and cell death pathways, we screened for caspase activity and expression of pro-survival proteins downstream of these kinases in cells proficient and deficient for SMNDC1. We found that PDAC and OvCa cells with acute loss of SMNDC1 had increased expression and activity of cleaved caspases and PARP (**Supplementary Fig. S3C**) and upregulation of pro-apoptosis members BIM, BAK and BAX of BCL-2 family (**Fig. 3F**). Consistently, SMNDC1 silencing led to downregulation of survival pathway members MCL1, BCL_XL_, and BCL_W_ of the BCL-2 family (**Fig. 3G**), which also contributes to the activation of cellular intrinsic apoptosis pathway. To validate that these results were not off-target effects of the shRNAs, we upregulated SMNDC1 and found that conversely, expression of BCL_XL_ was increased and BAX was decreased upon SMNDC1 overexpression (**Fig. 3H**). These results demonstrated that PDAC and OvCa cells harboring high expression of SMNDC1 display increased activity and expression of ERK1/2 and other proteins involved in cell proliferation and while decreased activity of pro-apoptotic pathways.

**Figure 3.**
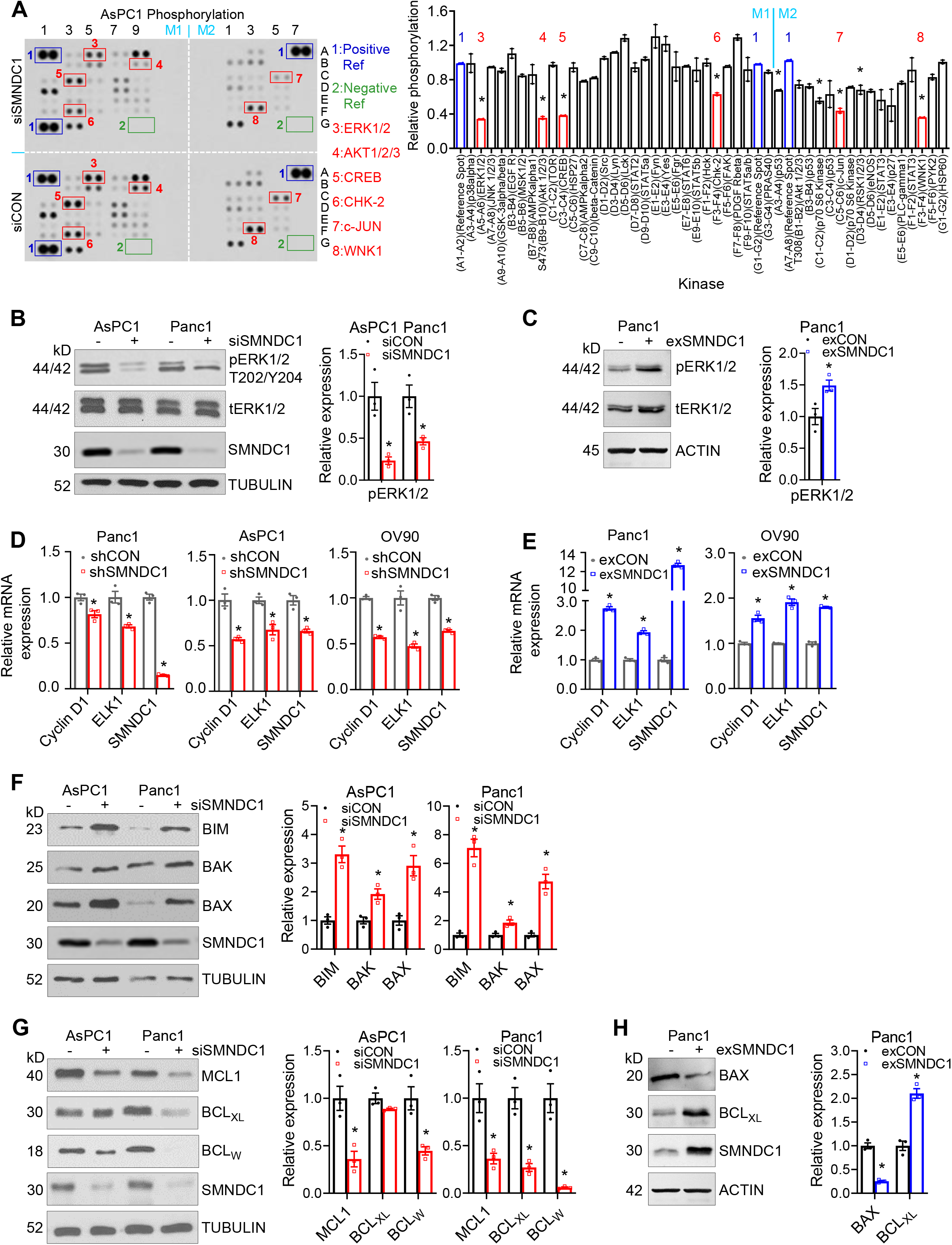
SMNDC1 upregulates the ERK pathway and blocks pro-apoptotic pathways. **A**, SMNDC1 silencing changed kinase phosphorylation. Seventy-two hours after siRNA transfection, AsPC1 cells were starved for 16 h and then stimulated with 10% FBS for 10 min. Cell lysates were incubated with membranes (M1, M2) printed with antibodies to kinase phosphorylation sites (Phospho-Kinase Array). Phosphorylation change was quantified by ratio of knockdown to control. Some significantly inhibited kinases by SMDC1 silencing were highlighted. **B**, ERK1/2 phosphorylation confirmation by western blot using AsPC1 and Panc1 cells. TUBULIN was used as a loading control. **C**, ERKk1/2 phosphorylation in cells with SMNDC1 stable overexpression. ex: overexpression. ACTIN was used as a loading control. **D, E**, mRNA expression of transcriptional target genes of ERK1/2 in SMNDC1 stable silence (**D**) or overexpression (**E**) cells. **F**, Expression of pro-apoptosis members in BCL-2 family in SMNDC1 knockdown cells and quantification of relative expression. **G**, Expression of anti-apoptosis/pro-survival members in BCL-2 family in SMNDC1 knockdown cells and quantification of relative expression. **H**, Expression of typical apoptosis/survival members at SMNDC1 stable overexpression. *, p<0.05.

### SMNDC1 promotes inclusion of guanine-rich cassette exons, impacting expression of ERK1

SMNDC1 was originally reported as a splicing regulator required for the assembly of the U4/U5/U6 tri-snRNP complex in the spliceosome (6-8), however, its role in specific splicing changes has not been reported. To determine the involvement of SMNDC1 in RNA splicing patterns in cancer, we performed deep RNA-seq and RNA splicing analyses in PDAC and OvCa cell lines with either expression or knockdown of SMNDC1. This identified differences in alternative RNA splicing of mainly cassette exon utilization in cells expressing SMNDC1 compared to cells with knockdown across the genome and having different coding and non-coding fates (**Fig. 4A-B, Supplementary Fig. S4A-F, Supplementary Tables ST2 and ST3**). Given that splicing changes in exons commonly occur in a sequence-specific manner, we characterized the nucleotide sequences enriched in retained versus skipped cassette exons in the context of SMNDC1, to understand the basis for alterations in cassette exon usage in tumors that overexpress SMNDC1. Exons retained in the presence of SMNDC1 in both PDAC and OvCa cells were rich in guanine, while skipped exons by SMDND1 were enriched in adenine/cytosine-rich sequences (**Fig. 4C**). These results link changes in RNA splicing of cassette exons to expression of SMNDC1. To prioritize the most relevant exonic changes induced by SMNDC1, we focused on identifying the common retained and skipped exons in PDAC and OvCa in the presence of SMNDC1. We found that among the common retained exons by SMNDC1, exon 4 of *MAPK3* (ERK1 protein) was among the most significant events, where ~50% of *MAPK3* transcripts had loss of exon 4 upon knockdown of SMNDC1 based on RNA-Seq (**Fig. 4D-E**). Exon 4 in *MAPK3*, is a 117bp exon, that encodes 39 residues in ERK1 (176-215aa), including the two most common phosphorylation residues that control ERK1’s activity (Thr202/Tyr204) and binding site to ERK2 (28) (**Fig. 4E**). We validated these differential *MAPK3* splicing differences by performing targeted PCRs in PDAC and OvCa lines with proficient and deficient SMNDC1 (**Fig. 4F-G**). Given that ERK1 and ERK2 are highly similar in sequence, we tested if SMNDC1 led to disrupted splicing of exon 4 of *MAPK1* (ERK2 protein), however, this was not the case (**Fig. 4H-I**). These results demonstrated that PDAC and OvCa cells harboring SMNDC1 display increased retention of exon 4 in *MAPK3* (ERK1).

**Figure 4.**
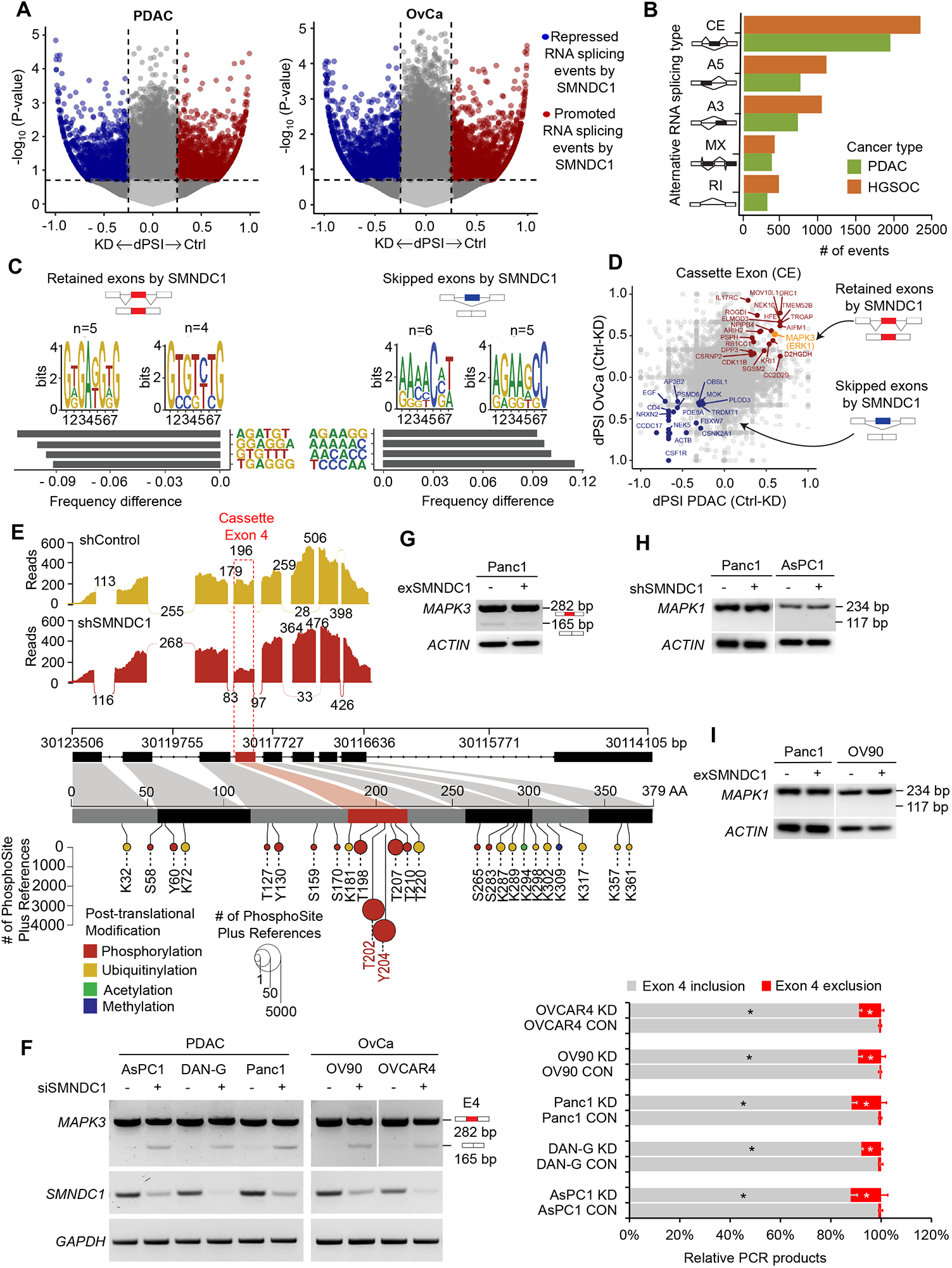
SMNDC1 promotes inclusion of guanine-rich cassette exons, impacting expression of ERK1. **A**, Repressed and promoted splicing events in PDAC and OvCa cells by SMNDC1. **B**, Alternative splicing types in PDAC and OvCa cells by SMNDC1. Here, cassette exon (CE) was detected as the most prominent splicing events in both cancer types. **C**, Kmer frequencies that are most up/downregulated between PDAC/OvCa shControl vs shSMNDC1 cells. **D**, Represents common CE that are either retained or skipped by SMNDC1 in both PDAC and OvCa cells. Exon 4 of MAPK3 (ERK1) considered significant event, as ~50% of MAPK3 transcripts had loss of exon 4 upon knockdown (KD) of SMNDC1. **E**, Sashimi plot (top) demonstrating skipping of *MAPK3* exon 4 (E4) upon KD of SMNDC1. The *MAPK3* transcript mapped to corresponding protein regions based on exon (bottom), annotated with reported post-translation modifications from the PhosphoSite database references. Hotspot phosphorylation residues T202 and Y204 are annotated in red. **F**, validation of CE of Exon 4 in *MAPK3* by performing targeted PCRs in PDAC and OvCa, deficient and, proficient (**G**) of SMNDC1. *MAPK3* E4 and CE of E4 (E4-) bands were quantified with ImageJ, adjusted by GAPDH control. Experiments were repeated 3 times. *, p<0.05, comparison of the same bands between CON and KD of the same cell lines. **H**, Targeted PCRs of deficient and, proficient (**I**) of SMNDC1 in PDAC and OvCa cells demonstrated no effect while targeting Exon 4 of MAPK1 (ERK2) which indicated its selectivity towards MAPK3.

### Inclusion of exon 4 in MAPK3 (ERK1) by SMNDC1 promotes cell proliferation and increased activity of ERK1

Deregulation of the EGFR-RAS-RAF-ERK ultimately leads to the activation of ERK1 and ERK2 by phosphorylation of the Thr202 and Tyr204 in ERK1 and to Thr183 and Tyr185 in ERK2. Given that overexpression of SMNDC1 in tumors promotes the retention of the exon encoding Thr202 and Tyr204 in ERK1, we first inquired about the relationship between SMNDC1 expression and mutations in the EGFR-RAS-RAF-ERK pathways, which ultimately lead to increased ERK1 activity. We performed genetic interaction analysis based on the concept of mutually exclusivity (29) which suggests that genetic alterations are exclusive to one another function within the same pathway. We found that mutations in EGFR-RAS-RAF, were mutually exclusive to overexpression of SMNDC1 (**Fig. 5A, Supplementary Fig. S5A**), suggesting the impact of SMNDC1 in the EGFR-RAS-RAF-ERK pathway and an independent mechanism by which non-mutant EGFR-RAS-RAF-ERK tumors, increase ERK 1 activity. We next sought to explore functional differences in ERK1 upon exon 4 (E4) inclusion and exclusion in PDAC cells. To test the role of exon 4 in ERK1 activity and conduct proof-of-principle experiments to validate the genetic exclusivity analyses, we designed a distinct morpholino antisense oligonucleotide (ASO) to block *MAPK3* E4 inclusion by blocking the 3’ splice sites (**Fig. 5B, Supplementary Fig. S5B**). The ASO was able to decrease the retention of the *MAPK3* E4 while maintaining constant the *MAPK3* mRNA levels relative to the control ASO (**Fig. 5C, Supplementary Fig. S5C**). We then tested the function of E4 in cell proliferation and cell signaling. First, we found that exclusion of E4 leads to decreased cell proliferation to almost the same extent as the knockdown of SMNDC1 (**Fig. 5D, Supplementary Fig. S5D**), suggesting the importance of E4 splicing regulation as one of the main mechanisms by which SMNDC1 leads to increased tumor growth. In addition, loss of E4 decreased the levels of phospho-ERK1 and expression of target transcriptional genes ELK1 and Cyclin D (**Fig. 5E-F, Supplementary Fig. S5E**). Furthermore, exclusion of E4 led to increased expression of BAX and decreased expression of BCL_XL_ (**Fig. 5G, Supplementary Fig. S5F**). Based on the critical function of E4 splicing in ERK1 function and signaling, we inquired if normal/benign somatic tissues and tumors from different anatomic sites control ERK1 activity by differentially regulating the inclusion or exclusion of this exon. Overall, we found that normal tissues tend to have less retention of E4 exon compared to tumors from different anatomical sites (**Fig. 5H**), and this was the case for pancreatic and ovarian malignant versus normal tissue (**Fig. 5I**). Furthermore, across normal organ tissues, we found differential retention of E4 ranging from appearing in only 25% of transcripts (liver) to almost 75% inclusion within transcripts (Testis, Ovary and others) (**Supplementary Fig. S5G**). On the other hand, in tumors we found that breast and kidney carcinomas incorporate E4 in almost 100% of *MAPK3* transcripts, while esophageal and stomach carcinomas only incorporate it in 50% of the transcripts (**Supplementary Fig. S5H**). These results suggest that differential splicing of E4 could lead to ERK1 signaling regulation in cells.

**Figure 5.**
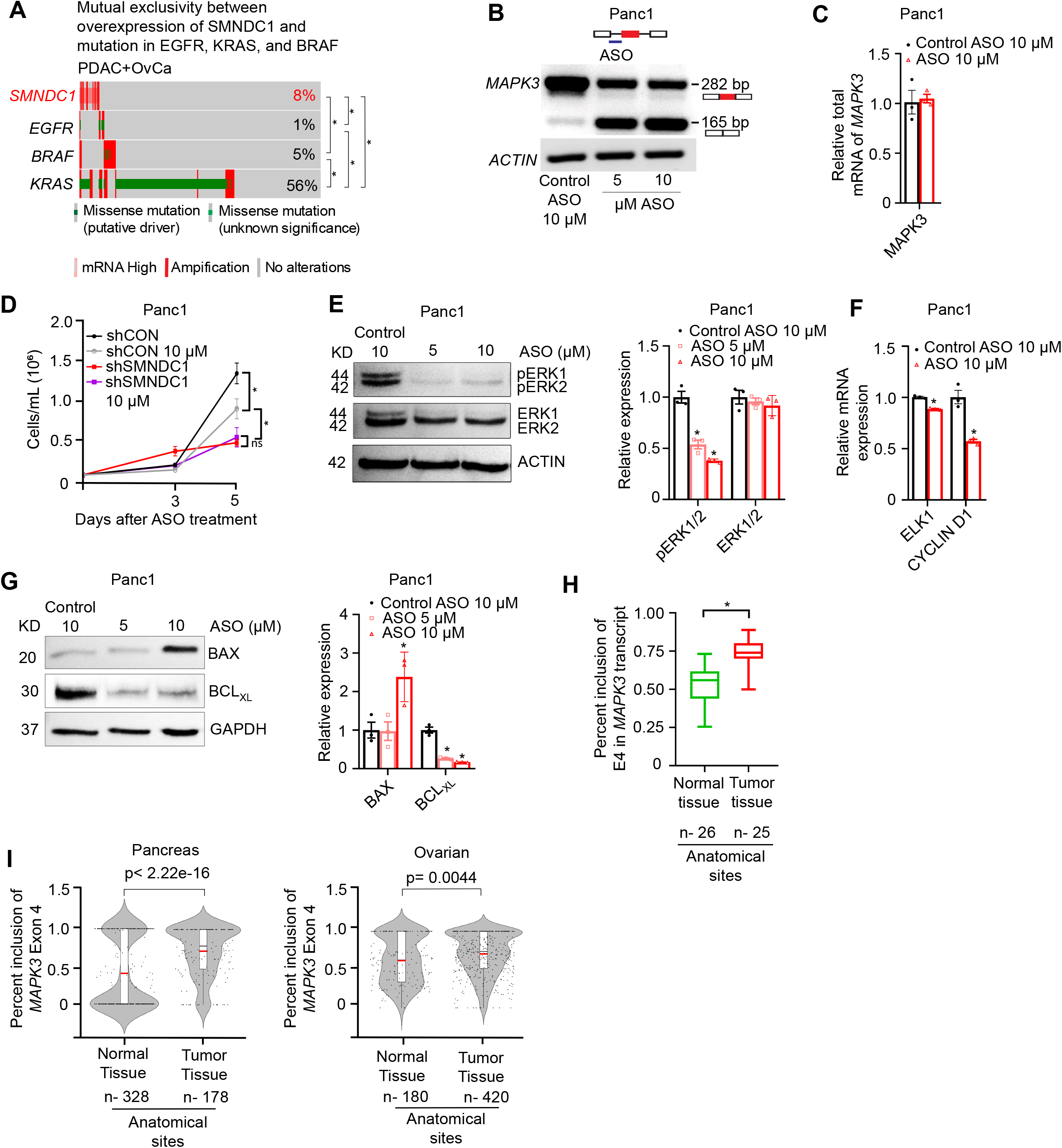
Inclusion of exon 4 of *MAPK3* (ERK1) by SMNDC1 promotes cell proliferation and increased ERK1 activity. **A**. Amplification and missense mutations in EGFR/KRAS/BRAF is mutually exclusive to the overexpression of SMNDC1 in PDAC+OvCa cohorts. **B**, Morpholino antisense oligonucleotides (ASOs) blocks 3’ splice site of MAPK3 E4 inclusion in Panc1 cells. **C**, Total RNA level for *MAPK3* after ASO treatment. **D**, Proliferation curve of Panc1 cells upon forced skipping of E4 in *MAPK3* alone or in combination with loss of SMNDC1 by stable silencing. **E**, Western blots for phospho-ERK1/2 and ERK1/2 upon skipping of E4 in *MAPK3* by ASO in Panc1 cells. **F**, mRNA expression of ELK 1 and Cyclin D1 upon forced skipping of E4 in *MAPK3*. **G**, Western blots for BAX (pro-apoptotic) and BCLXL (anti-apoptotic) upon skipping of E4 in *MAPK3*. **H**, Difference of mean difference in percentage spliced in (dPSI) of *MAPK3* Exon 4 between normal and tumor tissue across many anatomical sites. n-anatomical sites. **I**, Violin plot based on the percentage spliced in (PSI) value of MAPK3 Exon 4 of tumor and normal pancreatic and ovarian tissue. Here, Red bar indicate the mean PSI value. *, p<0.05

## DISCUSSION

Here we identify that overexpression of splicing factor SMNDC1 in tumors promotes their growth and survival by increasing ERK1 activation at the splicing level. Increased systematic sequencing of the genome and transcriptome of cancers has identified a variety of means by which RNA splicing is altered in cancer relative to normal cells. While studies have described how recurrent change-of-function mutations in splicing factors act as driver events in malignancy, less is known about the role of altered expression of splicing factors in oncogenic signaling and tumor growth. Here, we identify that overexpression of SMNDC1, which occurs in >10% of many carcinomas and is associated with poor patient outcome, leads to transcriptome-wide retention of sequence-specific exons in pancreatic and ovarian cancers that promote cell survival and anti-apoptotic signaling. Previous studies had described the role of SMNDC1 in the assembly of the spliceosome, however, our studies suggest that beyond this role, it promotes selective retention of exons which enhance oncogenic signaling. Our findings support other studies which have described that increased expression of splicing factors such as SRSF1 (30, 31), SRSF3 (32), and SRSF6 (33) acts as an oncogenic mechanism.

These results also highlight novel oncogenic mechanism of ERK1 activity regulation. ERK1 activation is fundamental for the development and progression of cancer. Since its discovery, studies have elucidated the effector role and activation mechanism of ERK1 downstream of the EGFR-RAS-RAF pathway (34). It is well established that phosphorylation at Thr202 and Tyr204 promotes the binding to ERK2 and nuclear translocation of ERK1, a process that is commonly deregulated in cancer by activating mutations in EGFR (7%), RAS (9%) and RAF (8%). However, we now demonstrate a non-genetic and plastic mechanism for ERK1 activation by overexpression of SMNDC1, where exon 4 of *MAPK3* (ERK1), which encodes both activation phosphorylation sites, is fully incorporated in all transcripts after RNA splicing, resulting in enhanced downstream signaling at the nucleus. This mechanism of kinase activity regulation by RNA splicing complements previous findings observed for CLK1 (35), and CHEK2 (36) kinases splicing and function. Since most common resistance mechanisms associated to targeted therapies against EGFR, RAS, RAF, MEK oncoproteins result in ERK reactivation, the targeting of ERK has become an emerging novel therapy in MAPK kinase-driven cancers. From a clinical standpoint our findings offer a novel mechanism and therapeutic strategy to target ERK in cancer by either splicing correcting approaches or by targeting SMNDC1 expression, which is specific to ovarian and pancreatic cancer cells.

To date, mutant EGFR/RAS/RAF/ERK pathway has been proven to be very difficult to target, preventing successful treatment of PDAC, OVCa and other carcinomas. Here, we identified that increased expression of splicing factor SMNDC1 modifies the splicing of exon 4 in *MAPK3* (ERK1) in a manner that promotes maximal activation of ERK1. Altering *MAPK3* exon 4 inclusion using antisense oligonucleotides significantly decreased oncogenic ERK1, survival and pro-apoptotic signaling resulting in decreased tumor cell growth. Collectively, these findings uncover a biologically critical and therapeutically exploitable mechanism to target carcinomas with increased SMNDC1 expression and ERK activity, and the basis for novel biomarker-driven treatment for PDAC and OvCa patients based on modulation of RNA splicing.

## AUTHORS’ DISCLOSURES

No disclosures were reported.

## AUTHORS’ CONTRIBUTIONS

**Y. Zhang:** Conceptualization, methodology, investigation, visualization, validation, formal analysis, writing–original draft, writing–review and editing. **M. A. Siraj:** Conceptualization, methodology, investigation, visualization, validation, formal analysis, writing–original draft, writing–review and editing. **P. Chakraborty:** Conceptualization, methodology, investigation, visualization, validation, formal analysis. **R. Tseng:** Methodology, formal analysis, writing–original draft. **L. Ku:** Methodology, investigation, validation. **S. Das:** Methodology, investigation, validation. **A. Dey:** Methodology, investigation, validation. **S. Dwivedi:** Methodology, investigation, validation. **G. Rao:** Methodology, investigation, validation. **M. Zhang:** Methodology, formal analysis, writing–original draft. **D. Yang:** Resources, supervision, funding acquisition, methodology, writing–review and editing. **M. N. Hossen:** Formal analysis. **W. Ding:** Formal analysis, methodology, resources. **K. Fung:** Visualization, formal analysis, resources. **R. Bhattacharya:** Conceptualization, formal analysis, methodology, resources. **L. Escobar-Hoyos:** Conceptualization, resources, supervision, funding acquisition, methodology, writing–original draft, writing–review and editing. **P. Mukherjee:** Conceptualization, resources, supervision, funding acquisition, methodology, writing–original draft, writing–review and editing.

## ABBREVIATIONS

A3: alternative 3’ splice site
A5: alternative 5’ splice site
AF: alternative first exon
AL: alternative last exon
AS: alternative splicing
BLCA: Bladder Urothelial Carcinoma
BRCA: Breast Invasive Carcinoma
CESC: Cervical Squamous Cell Carcinoma and Endocervical Adenocarcinoma
COAD: Colon Adenocarcinoma
EOC: epithelial ovarian cancer
ERK1: Extracellular Signal-Regulated Kinase 1
ESCA: Esophageal Carcinoma
GBM: Glioblastoma multiforme
GNP: gold nanoparticles
HGSOC: High Grade Serous Ovarian Cancer
HNSC: Head and Neck Squamous Cell Carcinoma
KIRC: Kidney Renal Clear Cell Carcinoma
KIRP: Kidney Renal Papillary Cell Carcinoma
LGG: Brain Lower Grade Glioma
LIHC: Liver Hepatocellular Carcinoma
LUAD: Lung Adenocarcinoma
LUSC: Lung Squamous Cell Carcinoma
MAPK3: Mitogen Activated Protein Kinase 3
MX: mutually exclusive exons
NAT: normal adjacent tissue
OVCa: Ovarian cancer
PAAD: Pancreatic Adenocarcinoma
PCPG: Pheochromocytoma and Paraganglioma
PDAC: pancreatic ductal adenocarcinoma
PRAD: Prostate Adenocarcinoma
RI: retained intron
SE: skipping exon
SARC: Sarcoma
SKCM: Skin Cutaneous Melanoma
SMNDC1: survival motor neuron domain containing 1
snRNP: small nuclear ribonucleoprotein
STAD: Stomach Adenocarcinoma
TMA: tissue microarray
TGCT: Testicular Germ Cell Tumors
THYM: Thymoma
THCA: Thyroid Carcinoma
TUNEL: terminal deoxynucleotidyl transferase dUTP nick end labeling
UCEC: Uterine Corpus Endometrial Carcinoma

## ACKNOWLEDGEMENTS

The results in Supplementary FigS1C are based upon data generated by the TCGA Research Network: https://www.cancer.gov/tcga. This work was supported by R01CA213278, R01CA136494, R01CA253391, R01CA260449 and P30 CA225500 Team Science grants (P.M.), the NCI K99-R00 CA226342-01 (to L.E-H), the William Raveis Charitable Fund / Rachleff Innovator of the Damon Runyon Cancer Research Foundation (to L.E-H), and the AACR Career Development Award to Further Diversity, Equity, and Inclusion in Pancreatic Cancer Research, (to L.E-H). Preparation of this publication was supported in part by the National Cancer Institute Cancer Center Support Grant P30CA225500 awarded to the University of Oklahoma Stephenson Cancer Center and an award from the Oklahoma Tobacco Settlement Endowment Trust to the University of Oklahoma Stephenson Cancer Center; Services from the Biostatistics and Research Design Shared Resource and the Office of Cancer Research were used. We also thank the Peggy and Charles Stephenson Cancer Center at the University of Oklahoma Health Sciences Center for a seed grant as well as thanking an Institutional Development Award (IDeA) Award from the National Institute of General Medical Sciences of the NIH (P20 GM103639) for the use of the Histology and Immunohistochemistry Core, which provided immunohistochemistry and image analysis services.

## FIGURE LEGENDS

**Supplementary Figure 1**. SMNDC1 is enriched by gold nanoparticles (GNP) and associated with cancer. **A**, Cell lysates from AsPC1 and Panc1 cells were incubated with GNP (40 μg/mL). GNPs binding proteins were boiled off and loaded for WB. 25 μg original lysate proteins were used as Input controls. ACTIN, TUBULIN and HSP90 were used as loading and binding controls. **B**. Oncoprint visualizing SMNDC1 genetic alteration in multiple cancer types. n: case number. **C**, Association of SMNDC1 mRNA expression with the survival of patients with cancer of liver, breast, head and neck, kidney or lung based on TCGA database, and ovary based on another GEO dataset (GSE32062). **D**, Intensity definition of TMA staining. Arrow heads: cancer cell nucleus (red). Scale bar: 100 µm. **E**, TMA H-score for all types of PDAC combined. n, case number. **F**, Association of tumor grade with patient survival based on PDAC TMA clinical data. Number in parentheses: case number. *, p<0.05.

**Supplementary Figure 2**. SMNDC1 promotes cancer cell proliferation, colony formation and tumor growth in mice. **A**, Efficiency of SMNDC1 knockdown by siRNA and shRNA. shRNAs targeting two different sequences of SMNDC1, designated as shSMNDC1-6 and shSMNDC1-7, were used. ACTIN was used as a loading control. **B**, Proliferation of cancer cells upon transient silencing of SMNDC1. **C**, Proliferation of cancer cells with stable silencing of SMNDC1. **D**, 3D colony formation (colony size) of cells upon transient silencing of SMNDC1. **E, F, G**, Example images, colony number and colony size of 3D colony formation assay using stable silencing cells. **H, I**, AsPC1 and OV90 xenograft tumor weight upon experiment termination. **J**, TUNEL staining of AsPC1 and OV90 xenograft tumors and quantification. Scale bar: 100 µm. *, p<0.05.

**Supplementary Figure 3**. SMNDC1 upregulates the ERK pathway and blocks pro-apoptotic pathways. **A**, Cell apoptosis, measured by caspase3/7 activity, and survival, measured by glycyl-phenylalanyl-aminofluorocoumarin (GF-AFC) live-cell protease activity, 96 h after siSMNDC1 transfection. Efficiency of SMNDC1 silencing was shown on the left. **B**, Phosphorylation confirmation of chosen kinases based on phospho-kinase array. **C**, Cleavage of caspases and PARP in common intrinsic apoptosis pathway in SMNDC1 knockdown cells and quantification of relative expression. *, p<0.05.

**Supplementary Figure 4**. SMNDC1 promotes inclusion of guanine/cytosine-rich cassette exons, impacting expression of ERK1. **A**, RNA quality for AsPC1 and (**B**) OV90 cells and corresponding SMNDC1 mRNA level after transient knockdown of SMNDC1. **C**, Volcano plot represent the repressed (Blue) and promoted (Red) splicing events in PDAC and OvCa by SMNDC1. Here, KD, SMNDC1 Knockdown cell lines, Ctrl, Control cell lines, dPSI, difference in percentage spliced in. **D**, Circus plots of PDAC and OvCa (**E**) demonstrate the up and down regulated cassette exon (CE) events in respective chromosomes positions. Here, each bubble represents the CE event, whereas the color represents the dPSI, and the size of the bubble represents the significance of the CE event. **F**, Meta-exon plot showing probability of secondary structure formation significantly up/down-regulated CE events categorized by their exon fates: SMNDC1 preferentially recognizes and retains 3’-UTR exons with higher probability of secondary structure formation. p(paired) describes the probability of a given base along the exon sequence to base-pair with another base, predicted by RNAfold at biological temperature (37º C). *, p<0.05.

**Supplementary Figure 5**. Inclusion of exon 4 (E4) of MAPK3 (ERK1) by SMNDC1 promotes cell proliferation and increased activity of ERK1 in AsPC1 cells. **A**. Amplification and missense mutations in EGFR/KRAS/BRAF is mutually exclusive to the overexpression of SMNDC1 in Liver+Stomach cohorts. **B**, Morpholino antisense oligonucleotides (ASOs) block *MAPK3* E4 inclusion in AsPC1 cells. **C**, Total RNA level for *MAPK3* after ASO treatment. **D**, Proliferation curve of ASPc1 cells with and, without ASOs. **E**, Western blots for phospho-ERK1/2, ERK1/2, (**F**) BAX (pro-apoptotic) and BCL_XL_ (anti-apoptotic) upon skipping of E4 in *MAPK3* by ASOs in AsPC1 cells. **G**, Percent spliced in (PSI) value of MAPK3 Exon 4 on normal tissue and (**H)** tumor tissue. *, p<0.05

## TABLES

**Supplementary Table 1 (ST1).** Patient and tissue specimen characteristics

|  | PDAC | Adenosquamous carcinoma | Mucinous cystadenoma of canceration | Total (100 cases) |
| --- | --- | --- | --- | --- |
| Gender |  |  |  |  |
| Male | 58 | 4 | 1 | 63 |
| Female | 35 | 2 |  | 37 |
| Age (years) | 61.6 | 59.5 | 83 |  |
| Grade |  |  |  |  |
| I-II | 10 |  | 1 | 11 |
| II | 53 | 4 |  | 57 |
| I-III | 1 |  |  | 1 |
| II-III | 20 | 1 |  | 21 |
| III | 8 | 1 |  | 9 |
| III-IV | 1 |  |  | 1 |
| Stage |  |  |  |  |
| 1A | 3 |  |  | 3 |
| 1B | 34 | 2 | 1 | 37 |
| 1-2 | 4 |  |  | 4 |
| 2A | 11 |  |  | 11 |
| 2 | 3 |  |  | 3 |
| 2B | 35 | 4 |  | 39 |
| 4 | 2 |  |  | 2 |
| Unknown | 1 |  |  | 1 |
| Survival status | Surgery date 2004.9-2011.12, visit date 2011.12. Being followed up for 3-7 years |  |  |  |
| survival | 25 (> 36 months) | 1 (> 47 months) | 1 (> 80 months) | 27 |
| deceased | 68 (< 43 months) | 5 (< 10 months) |  | 73 |

**Supplementary Table 2 (ST2).** Differential splicing event counts and distributions. dPSI>0.25, p<0.1

|  | A3 | A5 | AF | AL | MX | RI | SE | Total |
| --- | --- | --- | --- | --- | --- | --- | --- | --- |
| AsPC1 | 152 | 109 | 324 | 217 | 69 | 63 | 275 | 1209 |
| OV90 | 184 | 192 | 507 | 255 | 116 | 96 | 317 | 1667 |
| Common | 11 | 6 | 6 | 10 | 5 | 8 | 14 | 60 |

**Supplementary Table 3 (ST3).** Common cassette exon event. dPSI>0.25, p<0.1

| No | Gene | AsPC1 dPSI<br>KD-CON | p | OV90 dPSI<br>KD-CON | p |
| --- | --- | --- | --- | --- | --- |
| 1 | MAPK3 | 0.602 | 0.039 | 0.518 | 0.062 |
| 2 | DRC1 (1) | 0.667 | 0.099 | 0.774 | 0.083 |
| 3 | DRC1 (2) | 0.471 | 0.099 | 0.449 | 0.091 |
| 4 | PSPH | 0.308 | 0.062 | 0.514 | 0.001 |
| 5 | MOB1B | 0.27 | 0.056 | 0.316 | 0.026 |
| 6 | THAP9-AS1 | 0.295 | 0.094 | 0.335 | 0 |
| 7 | INO80C | 0.408 | 0.032 | 0.425 | 0.016 |
| 8 | RPL17-C18orf32 | 0.46 | 0.009 | 0.268 | 0.013 |
| 9 | CDK11B | 0.325 | 0.007 | 0.272 | 0.004 |
| 10 | SLMAP | 0.272 | 0.064 | 0.287 | 0.039 |
| 11 | LRIT3 | -0.667 | 0.067 | -0.333 | 0.088 |
| 12 | CCDC17 | -0.667 | 0.071 | -0.516 | 0.082 |
| 13 | HSD17B13 | -0.666 | 0.062 | -0.666 | 0.056 |
| 14 | NAT1 | -0.316 | 0.09 | -0.607 | 0.055 |

**Supplementary Table 4 (ST4).** Antibodies used for western blotting

|  | Antibody | Dilution | Catalog | Provider |
| --- | --- | --- | --- | --- |
| 1 | Rabbit anti-Actin | 1:10000 | A2066 | Sigma-Aldrich, St. Louis, MO |
| 2 | Rabbit anti-Akt | 1:1000 | #9272 | Cell Signaling, Danvers, MA |
| 3 | Rabbit anti-Bak | 1:1000 | #12105 | Cell Signaling |
| 4 | Rabbit anti-Bax | 1:1000 | #5023 | Cell Signaling |
| 5 | Rabbit anti-Bcl<sub>w</sub> | 1:1000 | #2724 | Cell Signaling |
| 6 | Rabbit anti-Bcl<sub>XL</sub> | 1:1000 | #2764 | Cell Signaling |
| 7 | Rabbit anti-Bim | 1:1000 | #2933 | Cell Signaling |
| 8 | Rabbit anti-Cleaved Caspase 3 | 1:500 | #9664 | Cell Signaling |
| 9 | Rabbit anti-Cleaved Caspase 7 | 1:500 | #8438 | Cell Signaling |
| 10 | Rabbit anti-Cleaved Caspase 9 | 1:500 | #7237 | Cell Signaling |
| 11 | Rabbit anti-Cleaved PARP | 1:500 | #5625 | Cell Signaling |
| 12 | Rabbit anti-Erk1/2 | 1:1000 | #9102 | Cell Signaling |
| 13 | Rabbit anti-Mcl1 | 1:1000 | #94296 | Cell Signaling |
| 14 | Rabbit anti-Phospho-Akt (Thr308) | 1:1000 | #3038 | Cell Signaling |
| 15 | Rabbit anti-Phospho-Akt (Ser473) | 1:1000 | #4060 | Cell Signaling |
| 16 | Rabbit anti-Phospho- Erk1/2<br>(Thr202/Tyr204) | 1:1000 | #8544 | Cell Signaling |
| 17 | Rabbit anti-SMND C1 | 1:1000 | NBP2-20424 | Novus Biologicals, Littleton,<br>USA |
| 18 | Mouse anti-Tubulin | 1:5000 | #66031 | Proteintech, Rosemont, IL |
| 19 | Mouse anti-HSP90 | 1:10000 | ab13492 | Abcam |
| 20 | Goat anti-Rabbit-IgG | 1:10000 | A6154 | Sigma-Aldrich |
| 21 | Goat anti-Mouse-IgG | 1:10000 | A4416 | Sigma-Aldrich |

## METHODS

### Cell Culture

Human PaC cell lines AsPC1, BxPC3, HPAF2, MiaPaCa2, Panc04.03 and Panc1 (37, 38) were purchased from American Type Culture Collection (ATCC, Manassas, VA); DAN-G (39) was obtained from German Collection of Microorganisms and Cell Cultures (DSMZ, Braunschweig, Germany). Immortalized human normal pancreatic ductal epithelial cell line HPDEC was got from AddexBio (San Diego, CA). Human OvCa cell line A2780 and A2780-CP20 (here as CP20) were kind gifts from Dr. A Sood (MD Anderson Cancer Center, Houston, TX) (40); OV202 was a kind gift from Dr. C Conover (Mayo Clinic, Rochester, MN) (41); OVCAR2 and A1847 were kind gifts from Dr. S Johnson (University of Pennsylvania Cancer Center, Philadelphia, PA) (42); OVCAR3 and OV90 were from ATCC; OVCAR4 was from Sigma-Aldrich (St. Louis, MO). The SV40 transformed primary normal ovarian surface epithelial cell line OSE was a kind gift from Dr. V Shridhar (Mayo Clinic, Rochester, MN) (43). Human embryonic kidney 293T cell line (293T) was from ATCC. AsPC1, BxPC3, DAN-G, A1847, A2780, CP20, OV90, OVCAR2, OVCAR3 and OVCAR4 were grown in RPMI 1640 (10-040-CV, Corning, NY) supplemented with 10% Fetal Bovine Serum (FBS, 16000-044, Life technologies, Carlsbad, CA) and 1% Penn-Strep (15140-122, Life technologies). Panc04.03 was culture in RPMI 1640 supplemented with 15% FBS, 10 μg/mL Insulin (I9278, Sigma) and 1% Penn-Strep. HPAF2, Panc1 and 293T were propagated in DMEM (10-013-CV, Corning) supplemented with 10% FBS and 1% Penn-Strep. MiaPaCa2 was cultivated in DMEM supplemented with 10% FBS, 2.5% horse serum (26050070, Life technologies) and 1% Penn-Strep. HPDEC was fostered in Keratinocyte-SFM (17005042, Invitrogen, Carlsbad, CA) with human recombinant Epidermal Growth Factor and Bovine Pituitary Extract. OV202 was grown in MEMα (41061, ThermoFisher Scientific, Waltham, MA) with 10 % FBS and 1 % Penn-Strep. OSE was cultured in 1:1 MCDB105 and Medium199 (Corning, Corning, NY, USA) with 10%FBS and 10ng/ml EGF. All the cells were maintained at 37 °C in humidified 95% air and 5% CO_2_ atmosphere.

### Gene Knockdown and overexpression

For transient knockdown of SMNDC1 gene, 100 nM siRNA targeting SMNDC1 (SASI_Hs01 00210371, Sigma), or negative control siRNA (SIC001, Sigma), was transfected into 60-70% confluent cells grown on 60 mm culture dish (1156F05, Thomas Scientific, Swedesboro, NJ) with 12 μl transfection reagent Lipofectamine RNAiMAX (13778150, Invitrogen) in 2.2 ml volume. For stable knockdown of SMNDC1 gene, 0.5 μg TRC2-pLKO-puro vector encoding shRNA (TRCN0000377256-NM_005871.3-637s21c1, designated here as shSMNDC1-6, or TRCN0000369078-NM_005871.3-724s21c1, designated here as shSMNDC1-7, Sigma) targeting different sequences of SMNDC1, or negative control shRNA (TRC2 pLKO.5-puro Non-Target shRNA Control Plasmid DNA, SHC216, Sigma) were transfected to 60-70% confluent 293T cells grown on 6-well plates (1156D98, Thomas Scientific, Swedesboro, NJ) with 4.6 μl Lentiviral Packaging Mix (SHP001, Sigma) and 2.7 μl FuGENE 6 Transfection Reagent (E2691, Promega, Madison, WI) in 1.5 ml volume. Lentiviral supernatants were collected 2 and 3 days after transfection, filtered with 0.2 μm filters (4612, Pall Corporation, New York, NY) and added to target cells on 6-well plates. Target cells were selected with 2 μg/ml puromycin (A1113803, ThermoFisher) for 2 weeks commencing 48 h after infection to establish stable lines. Stable knockdown cells were maintained in complete medium with 1 μg/ml puromycin (44). For stable overexpression of SMNDC1, 1μg pCMV3-SMNDC1-C-FLAG (HG14294-CF, Sino Biological US Inc, Houston, TX) or pCMV3-C-FLAG-EMPTY was transfected to cells in 6-well plate. The cells were then selected and maintained in 200 μg/ml hygromycin (10843555001, Sigma).

### Proliferation Assays

Cell proliferation was evaluated by manual cell counting and CyQuant assay. For manual cell counting, 1×10^4^ cells were seeded onto each well in 24-well plates (1156F00, Thomas Scientific). Cells were counted daily 2 days later for 4 days with trypan blue exclusion method. For CyQuant assay, 2×10^3^ cells were seeded onto each well in 96-well plates (1156F02, Thomas Scientific). Cells were labeled with Cyquant NF dye reagent (C35006, ThermoFisher) for 30 min and the fluorescence intensity (excitation at 480 nm, emission at 520 nm) was measured using CLARIOstar microplate reader (BMG Labtech, Cary, NC) from the next day for 4-5 consecutive days.

### Animal Studies

Female athymic nude mice (NCr-nu; 5-week old, Harlan Laboratories, Indianapolis, IN) were housed and maintained under pathogen-free conditions in facilities approved by American Association for Accreditation of Laboratory Animal Care and in accordance with regulations and standards of US Department of Agriculture, Department of Health and Human Services, and National Institutes of Health. Studies were approved and supervised by University of Oklahoma Health Science Center Institutional Animal Care and Use Committee. SMNDC1 stable knockdown and control AsPC1 or OV90 cells (2×10^6^ cells/100ul PBS) were inoculated subcutaneously (SC) to the flank of the mice (n=9 for AsPC1, n=10 for OV90). Mice weights were recorded weekly, and their health and behavior were monitored daily. Tumor sizes were measured with caliper twice a week (AsPC1) or every two days (OV90) starting from 9 days (AsPC1) or 6 days (OV90) after implantation. Tumor volume was calculated as V = W^2^ × L/2, where W is tumor width and L is tumor length. Experiment was terminated 37 days (AsPC1) or 16 days (OV90) after cell implantation. Tumors were measured (volume and weight), photographed, and preserved in 10% formalin solution (HT501128, Sigma) or -80°C.

### Immunohistochemical stains

Tumor grafts were fixed in 10% formalin solution for 24 h, transferred to 70% ethanol, embedded with paraffin, serially sectioned at 4 μm thickness, and mounted on positively charged slides. Human PaC tissue microarrays (TMA) HPan-Ade180Sur-02 was purchased from US Biomax (Rockville, MD). The sections or TMAs were incubated at 60°C for 45 min, followed by deparaffinization and rehydration in an automated Multistainer (ST5020, Leica, Wetzlar, Germany). The slides were then transferred to the BOND-III automated IHC Stainer (Leica) for stepwise incubation with BOND Epitope Retrieval Solution 1 pH 6 (AR9961, Leica) at 100°C for 20 min, 5% goat serum (01-6201, ThermoFisher) at 25°C for 30 min, Peroxidase Block (RE7101, Leica) at 25°C for 10 min, and primary antibodies rabbit anti-SMNDC1 (1:1500, NBP2-20424, Novus Biologicals) or rabbit anti-Ki67 (1:1000) at 4°C for 16 h. The Bond Polymer Refine Detection System (DS 9800, Leica) was then used to localize primary antibodies, visualize the targets, and counterstain cell nuclei according to the manufacturer’s instructions. The slides were dehydrated in Multistainer and mounted with MM 24 Mounting Media (3801120, Leica). To detect apoptosis in tissue, terminal deoxynucleotidyl transferase dUTP nick end labeling (TUNEL) was performed using an In Situ Cell Death Detection Kit, AP (11684809910, Sigma). Briefly, after deparaffinization and rehydration, the sections were subjected to cell permeabilization with 0.1M citrate buffer, pH 6.0, non-specific reaction blocking with 10% FBS + 3% BSA in PBS at 37°C for 20 min, incubation with 50 μl TUNEL mixture at 37°C for 1 h, followed by reaction in 50 μl converter AP solution for 30 min. The cells were illustrated with Fast Red Substrate (HK182-5KE, BioGenex, Fremont, CA) and counterstained with hematoxylin. The stained slides were mounted with Aquamount (13800, ThermoFisher). Negative (omission of primary antibody) control was stained parallelly and no staining was observed under these conditions (45). Whole slides were scanned using Aperio eSlide Manager (Leica). H-score was estimated by two independent pathologists without knowledge of patient clinical outcome based on the formula H=0 × (% cells 0+) + 1 × (% cells 1+) + 2 × (% cells 2+) + 3 × (% cells 3+), where the staining intensity of 0=negative, 1=weak, 2=moderate and 3=strong. The two H-scores for each spot were then averaged (46, 47). For TUNEL assay analysis, staining was converted into gray images and the intensity for these images was quantified by ImageJ (National Institutes of Health, Rockville, MD), where background intensity was nullified. Two PDAC cases whose tissue spots were detached by the staining and one PDAC case without staging information were removed from the analysis. The staining was then scored by two independent pathologists based on H-score estimation, which takes into account both intensity and percentage of the epithelial cells staining (33, 46, 47).

We analyzed PaCTMA data to see whether SMNDC1 expression is associated with tumor grade and patient survival. To do this, we grouped PDAC survival data according to available grades (the one grade I-III case was grouped to II-III, the one III-VI case was removed as the tissue spot was detached, **Supplementary Table ST1**).

### Western blotting

For gene transient knockdown cells, cell lysates were collected 3 or 4 days after transfection. For protein phosphorylation detection, cells were starved in serum-free medium for 24 h and then incubated with complete medium for 10 min before cell collection. For preparation of cell lysate, medium was removed, and cells were placed on ice, washed twice with ice-cold PBS and lysed with RIPA buffer (BP-115, Boston BioProducts, Ashland, MA) containing Protease and Phosphatase Inhibitor Cocktail (78440, ThermoFisher). Protein concentration of the lysates was measured using BCA Protein Assay Kit (23227, ThermoFisher). Lysates (10-100 μg proteins/lane) were loaded to and separated on 6%-15% SDS-PAGE gels and electrophoretically transferred to PVDF membrane (1620177, Bio-Rad, Hercules, CA). The membranes were blocked in 4% BSA/PBST (0.1% Tween-20, P1379, Sigma-Aldrich, in PBS) at room temperature for 1 h, incubated with primary antibodies at 4°C for 16 h, followed by washing and incubation with HRP-coupled secondary antibodies for 1 h at ambient temperature. Signals were visualized with SuperSignal West Femto (TI271896A, ThermoFisher) or Clarity Western ECL Substrates (1705061, Bio-Rad). Blots were scanned with Color LaserJet Pro MFO M477fdn (HP, Palo Alto, CA). Primary and secondary antibodies used are listed in **Supplementary Table ST4**.

For ASO transfected PDAC cells, protein lysates were collected 96 h after morpholino delivery. Cell lysate was prepared with Pierce™ RIPA buffer (89900, Thermo Scientific, Rockford, IL) containing Protease and Phosphatase Inhibitor Cocktail (1861281, Thermo Scientific, Rockford, IL) after washing cell twice with ice-cold PBS. Protein concentration of the lysates was measured using Quick Strat Bradford assay (5000205, Bio-Rad, Hercules, CA). Lysates (32 μg proteins/lane) were loaded to and separated on 10% SDS-PAGE gels and electrophoretically transferred to PVDF membrane (1620264, Bio-Rad, Hercules, CA). The membranes were blocked in 5% BSA/TBST (0.1% Tween-20, P1379, Sigma-Aldrich, in PBS) at room temperature for 1 h, incubated with primary antibodies at 4°C for 24 h, followed by washing and incubation with HRP-Conjugated secondary antibodies for 1 h at ambient temperature. Signals were visualized with Radiance Plus (AC2103, AZURE, Dublin, CA). Blots were scanned with iBright imaging system (Invitrogen).

### Caspase 3/7 Activity Assay and Cell Viability

Caspase 3/7 activity and cell viability were determined by ApoTox-Glo assay kit (G6321, Promega). Briefly, SMNDC1 silencing or control cells were seeded to 96-well plate 1000 cells/well 1 day after siRNA transfection, and cell apoptosis or viability was measured 3 days after using a luminogenic caspase3/caspase7 substrate the tetra-peptide sequence DEVD or using GF-AFC live-cell protease (48).

### Antibody Array Assay

Seventy-two hours after siRNA transfection, AsPC1 cells were starved for 16 h and then stimulated with 10% FBS for 10 min. Cells were collected and lysed. 300 μg cell lysates were used for phosphorylation analysis with Human Phospho-Kinase Array Kit (ARY003B, R&D Systems, Minneapolis, MN) following the manufacturer’s instruction.

### RNA Isolation and Sequencing

Total RNA from AsPC1 or OV90 cells transfected with 100 nM siSMNDC1 or siCON for 48 h was extracted using Quick-RNA MiniPrep (R1055, Zymo Research, Irvine, CA). Three biological repeats were performed. RNA sequencing was performed with an Illumina Hiseq 2000 by the Oklahoma Medical Research Foundation (OMRF) Sequencing Core.

### RT-PCR and qPCR

cDNA was synthesized using iScript Select cDNA Synthesis Kit (Bio-Rad) with “oligo(dT) + Random” primers and 0.1-1 µg total RNA. MAPK3 and GAPDH PCR primers were designed using PrimerQuest Tool (IDT, Coralville, Iowa) and synthesized by IDT. MAPK3 primer sequences are: sense 5’GTGCTCCACCGAGATCTAAAG3’, antisense 5’GTTGAGCTGATCCAGGTAGTG3’. GAPDH primer sequences are: sense 5’GGTCGGAGTCAACGGATTT3’, antisense 5’TCTTGAGGCTGTTGTCATACTT3’. SMNDC1, ELK1, CYCLIN D1, MAPK1 and ACTIN PCR primers were designed using “Primer-Blast” Tool (NCBI) and synthesized by Keck Oligo Synthesis Resource at Yale. SMNDC1 primer sequences are: sense 5’TCCAGGGTGTGTTAACAAAAGGG3’, antisense 5’CTTGCATGAATGCCGTGTGC3’. ELK1 primer sequences are: sense 5’AGATCACCCAACCGCAGAAG3’, antisense 5’GGGTCAAGGTATGTGTGGGG3’. CYCLIN D1 primer sequences are: sense 5’AGAACAAGCAGACCATCCGC3’, antisense 5’GTCCTTGTTTAGCCAGAGGC3’. MAPK1 primer sequences are: sense 5’GAAGACACAACACCTCAGCA3’, antisense 5’CCAAAATGTGGTTCAGCTGG3’. ACTIN primer sequences are: sense 5’CACTGTCGAGTCGCGTCC3’, antisense 5’CGCAGCGATATCGTCATCCA3’. SMNDC1 primers (Hs_SMNDC1_1_SG QuantiTect Primer Assay QT00014035) were from Qiagen (Hilden, Germany). PCR was performed with Hot Start Taq Polymerase (M0495S, NEB, Ipswich, MA). PCR condition for MAPK3: 33-35 cyc, 95°C30s-50°C20s-68°C20s; for MAPK1: 50 cyc, 95°C30s-50°C20s-68°C20s; For SMNDC1: 30 cyc, 95°C30s-60°C20s-68°C20s; For GAPDH: 25 cyc, 95°C30s-60°C20s-68°C30s. PCR products were run in 2% or 4% agarose gel. qPCR was performed using iTaq SYBR Green (Bio-Rad) following supplier’s direction. Ten µL reaction volume including 1 µL cDNA were used.

### Nucleotide Motif Enrichment Analyses

Statistically enriched nucleotide motifs were identified with ab initio method MEME suite by querying all k-mers of length 4, 5 or 6. An enrichment statistic was defined for each k-mer as the occurrence of each k-mer in all cassette exons that were promoted or repressed in the SMNDC1 knockdown or control samples. Statistically significant enrichment was identified with the nonparametric Wilcoxon test, performed using the distributions of occurrence for each k-mer in promoted versus repressed exons. Differentially enriched motifs with an adjusted p value < 25% were considered significant. Consensus motifs were then identified by performing k-means clustering on significantly enriched or depleted motifs with n = 5 centers (49).

### RNA-seq alignment and splicing analysis

Deep RNA sequencing of Panc1 and OV90 WT and SMNDC1 knockdown (KD) samples (n = 3 each) were aligned through Kallisto against the hg38 GENCODE human reference files with 50 bootstrap samples and standard parameters for all other options. The raw transcript-per-million counts are then inputted into SUPPA2 to obtain percent-spliced-in (PSI) estimates for all hg38 annotated splicing events. The change in percent-spliced in (ΔPSI) in WT vs. SMNDC1 KD for each event was calculated by subtracting the mean PSI value of WT triplicates against KD triplicates. The p-value for each event was calculated via unpaired t-test. Differentially spliced events were obtained with a threshold of |dPSI| ≥ 0.25 and p-value ≤ 0.2.

### RNA secondary structure motif analysis

The HGNC gene symbol, Ensembl transcript and exon IDs, the gene, transcript, exon, UTR, and CDS start/end positions, transcript, and exon biotypes, as well as the exon number of each skipped exon (SE) event was obtained through querying the annotated Ensembl gene ID, exon coordinates, and sense against the Ensembl database via the R biomaRt package. The sequences of each SE event as well as 50bp of each up/downstream intron were obtained via the getSeq() function of the R BSgenome package, queried against the UCSC hg38 human genome sequence reference. The sequences of each event were then piped into the RNAfold 2.5.1 program with default parameters, predicting the probability-paired, or p(paired), of each base position based on simulated secondary structure folding of the given sequence. The meta-exon distribution of p(paired) for all SE events was then aggregated by linear interpolation of each SE event and averaging each relative base position. The meta-intron distributions flanking 50bp up/downstream were aggregated by averaging each absolute base position.

### Mutually exclusivity analysis

Mutually exclusivity is tested using the one-sided Fisher exact test. The p-value, implying the significance, is determined under the null hypothesis. The Log2Odds Ratio is determined by Log2(OR) = Log2[(*A* \**D*)/(*B* * *C*)], where A is the number of alterations in both genes; B is the number of alterations in gene1 but not gene2; C is the number of alterations in gene2 but not gene1; and D is the number of alterations in neither genes. Log2Odds Ratio < -1 implies the frequency of occurrence of alterations in two genes have tendency toward mutually exclusivity. - 1 < Log2Odds Ratio < 1 implies the frequency of occurrence of alterations in two genes have no association. Log2Odds Ratio > 1 implies the frequency of occurrence of alterations in two genes have tendency toward co-occurrence. For the total six raw datasets of Breast, Liver, LUAD, PDAC, Ovary, Stomach, 52 combinations of cancers were taken for testing the mutually exclusivity. The final mutually exclusivity results are selected by the cutoff of Log2Odds Ratio < - 1 with p-value < 0.05.

### In Vitro Anti-sense Oligonucleotide (ASO) Transfection

Manufacturer’s (GeneTools LLC) guidelines were followed to deliver morpholino into Panc-1 and AsPc1 cell lines. Briefly, cells were cultured in 6-well plate till to become ~ 90% confluent. Before using the morpholino, spent culture media was replace with fresh media supplemented with 10% FBS. A final concentration of 10 µM of morpholino oligo was added and swirl well to mix. A final concentration of 6 µM Endo-Porter was mixed after adding morpholino. ASO target sequence was GGCAGCCCCTCCTACCTTGGAGTTC. The predicted melting temperature (Tm) of the targeted oligo was optimized for use at 37°C compared to other designed oligos which had a lower predicted Tm. In addition, the selected oligo targeted to the donor splice junction which is preferable for specificity.

### Comparative analysis of the expression of MAPK3 exon 4 in normal tissue of different organ system

TCGA/GTEx data was analyzed to understand the comparative scenario of the expression of MAPK3 exon 4 in normal tissue of different organ system. Violin plots were developed based on the PSI value of MAPK3 exon 4 of each normal and tumor tissue sample. For both normal and tumor tissue, the organ type which demonstrated a sample size (N) >100 was considered for further analysis. In the figure, the horizontal bar in the box plot represents the median, whereas the red lines in the box plots of violin plots show the means. Both PDAC and OvCa between normal and tumor tissue were found statistically significant under t test while comparing the percent spliced in (PSI) values.

### Statistics

Parametric and non-parametric tests (t-test, Wilcoxon-Mann-Whitney) will be used to compare differences in tumor sizes, metastases, and phenotypes. Multiple comparison adjustments will be used when multiple variables are being assessed. For survival differences, we will use log-rank test and Cox’s proportional hazards model. Other statistics was accomplished by one-way ANOVA with Tukey’s post-test. GraphPad Prism 8 software was used. Data are expressed as mean ± SEM. P value ≤0.05 was considered statistically significant, except some situations specified in the context.

## Notes

### Competing Interest Statement

The authors have declared no competing interest.

## REFERENCES

1. S. C. Bonnal, I. Lopez-Oreja, J. Valcarcel, Roles and mechanisms of alternative splicing in cancer - implications for care. Nat Rev Clin Oncol 17, 457–474 (2020).

2. S. C. Lee, O. Abdel-Wahab, Therapeutic targeting of splicing in cancer. Nat Med 22, 976–986 (2016).

3. Y. Zhang, J. Qian, C. Gu, Y. Yang, Alternative splicing and cancer: a systematic review. Signal Transduct Target Ther 6, 78 (2021).

4. L. Escobar-Hoyos, K. Knorr, O. Abdel-Wahab, Aberrant RNA Splicing in Cancer. Annu Rev Cancer Biol 3, 167–185 (2019).

5. G. Neubauer et al., Mass spectrometry and EST-database searching allows characterization of the multi-protein spliceosome complex. Nat Genet 20, 46–50 (1998).

6. J. Rappsilber, P. Ajuh, A. I. Lamond, M. Mann, SPF30 is an essential human splicing factor required for assembly of the U4/U5/U6 tri-small nuclear ribonucleoprotein into the spliceosome. J Biol Chem 276, 31142–31150 (2001).

7. G. Meister et al., SMNrp is an essential pre-mRNA splicing factor required for the formation of the mature spliceosome. EMBO J 20, 2304–2314 (2001).

8. J. T. Little, M. S. Jurica, Splicing factor SPF30 bridges an interaction between the prespliceosome protein U2AF35 and tri-small nuclear ribonucleoprotein protein hPrp3. J Biol Chem 283, 8145–8152 (2008).

9. K. Talbot, I. Miguel-Aliaga, P. Mohaghegh, C. P. Ponting, K. E. Davies, Characterization of a gene encoding survival motor neuron (SMN)-related protein, a constituent of the spliceosome complex. Hum Mol Genet 7, 2149–2156 (1998).

10. S. Kilpinen et al., Systematic bioinformatic analysis of expression levels of 17,330 human genes across 9,783 samples from 175 types of healthy and pathological tissues. Genome Biol 9, R139 (2008).

11. A. Sveen, S. Kilpinen, A. Ruusulehto, R. A. Lothe, R. I. Skotheim, Aberrant RNA splicing in cancer; expression changes and driver mutations of splicing factor genes. Oncogene 35, 2413–2427 (2016).

12. Y. J. Guo et al., ERK/MAPK signalling pathway and tumorigenesis. Exp Ther Med 19, 1997–2007 (2020).

13. E. Martinelli, F. Morgillo, T. Troiani, F. Ciardiello, Cancer resistance to therapies against the EGFR-RAS-RAF pathway: The role of MEK. Cancer Treat Rev 53, 61–69 (2017).

14. N. Sugawa, A. J. Ekstrand, C. D. James, V. P. Collins, Identical splicing of aberrant epidermal growth factor receptor transcripts from amplified rearranged genes in human glioblastomas. Proc Natl Acad Sci U S A 87, 8602–8606 (1990).

15. U. H. Weidle, D. Maisel, S. Klostermann, E. H. Weiss, M. Schmitt, Differential splicing generates new transmembrane receptor and extracellular matrix-related targets for antibody-based therapy of cancer. Cancer Genomics Proteomics 8, 211–226 (2011).

16. H. Wang et al., Identification of an exon 4-deletion variant of epidermal growth factor receptor with increased metastasis-promoting capacity. Neoplasia 13, 461–471 (2011).

17. F. D. Tsai et al., K-Ras4A splice variant is widely expressed in cancer and uses a hybrid membrane-targeting motif. Proc Natl Acad Sci U S A 112, 779–784 (2015).

18. C. Papin, A. Denouel-Galy, D. Laugier, G. Calothy, A. Eychene, Modulation of kinase activity and oncogenic properties by alternative splicing reveals a novel regulatory mechanism for B-Raf. J Biol Chem 273, 24939–24947 (1998).

19. P. I. Poulikakos et al., RAF inhibitor resistance is mediated by dimerization of aberrantly spliced BRAF(V600E). Nature 480, 387–390 (2011).

20. G. Maik-Rachline, I. Wortzel, R. Seger, Alternative Splicing of MAPKs in the Regulation of Signaling Specificity. Cells 10, (2021).

21. T. G. Boulton et al., ERKs: a family of protein-serine/threonine kinases that are activated and tyrosine phosphorylated in response to insulin and NGF. Cell 65, 663–675 (1991).

22. K. Giri et al., Understanding protein-nanoparticle interaction: a new gateway to disease therapeutics. Bioconjug Chem 25, 1078–1090 (2014).

23. D. R. Rhodes et al., ONCOMINE: a cancer microarray database and integrated data- mining platform. Neoplasia 6, 1–6 (2004).

24. X. Deschenes-Simard, F. Kottakis, S. Meloche, G. Ferbeyre, ERKs in cancer: friends or foesã Cancer Res 74, 412–419 (2014).

25. K. B. Acosta, M. M. Tibolla, M. M. Tiscornia, M. A. Lorenzati, P. D. Zapata, Recent patents related to phosphorylation signaling pathway on cancer. Recent Pat DNA Gene Seq 5, 175–184 (2011).

26. C. G. Proud, Phosphorylation and Signal Transduction Pathways in Translational Control. Cold Spring Harb Perspect Biol 11, (2019).

27. M. Linder et al., EGFR is required for FOS-dependent bone tumor development via RSK2/CREB signaling. EMBO Mol Med 10, (2018).

28. I. Wortzel, R. Seger, The ERK Cascade: Distinct Functions within Various Subcellular Organelles. Genes & Cancer 2, 195–209 (2011).

29. J. Cisowski, M. O. Bergo, What makes oncogenes mutually exclusiveã Small GTPases 8, 187–192 (2017).

30. O. Anczukow et al., SRSF1-Regulated Alternative Splicing in Breast Cancer. Mol Cell 60, 105–117 (2015).

31. O. Anczukow et al., The splicing factor SRSF1 regulates apoptosis and proliferation to promote mammary epithelial cell transformation. Nat Struct Mol Biol 19, 220–228 (2012).

32. R. Jia, C. Li, J. P. McCoy, C. X. Deng, Z. M. Zheng, SRp20 is a proto-oncogene critical for cell proliferation and tumor induction and maintenance. Int J Biol Sci 6, 806–826 (2010).

33. M. Cohen-Eliav et al., The splicing factor SRSF6 is amplified and is an oncoprotein in lung and colon cancers. J Pathol 229, 630–639 (2013).

34. F. Marampon, C. Ciccarelli, B. M. Zani, Biological Rationale for Targeting MEK/ERK Pathways in Anti-Cancer Therapy and to Potentiate Tumour Responses to Radiation. Int J Mol Sci 20, (2019).

35. P. I. Duncan, D. F. Stojdl, R. M. Marius, J. C. Bell, In vivo regulation of alternative pre-mRNA splicing by the Clk1 protein kinase. Mol Cell Biol 17, 5996–6001 (1997).

36. V. Staalesen et al., Alternative splicing and mutation status of CHEK2 in stage III breast cancer. Oncogene 23, 8535–8544 (2004).

37. E. L. Deer et al., Phenotype and genotype of pancreatic cancer cell lines. Pancreas 39, 425–435 (2010).

38. Z. Zhuang et al., IL1 Receptor Antagonist Inhibits Pancreatic Cancer Growth by Abrogating NF-kappaB Activation. Clin Cancer Res 22, 1432–1444 (2016).

39. G. L. Razidlo et al., Targeting Pancreatic Cancer Metastasis by Inhibition of Vav1, a Driver of Tumor Cell Invasion. Cancer Res 75, 2907–2915 (2015).

40. M. Moreno-Smith et al., ATP11B mediates platinum resistance in ovarian cancer. J Clin Invest 123, 2119–2130 (2013).

41. K. R. Kalli et al., Functional insulin receptors on human epithelial ovarian carcinoma cells: implications for IGF-II mitogenic signaling. Endocrinology 143, 3259–3267 (2002).

42. D. Roberts et al., Identification of genes associated with platinum drug sensitivity and resistance in human ovarian cancer cells. Br J Cancer 92, 1149–1158 (2005).

43. J. Chien et al., A role for candidate tumor-suppressor gene TCEAL7 in the regulation of c-Myc activity, cyclin D1 levels and cellular transformation. Oncogene 27, 7223–7234 (2008).

44. Y. Zhang et al., Prostaglandin E2 receptor 4 mediates renal cell carcinoma intravasation and metastasis. Cancer Lett 391, 50–58 (2017).

45. S. A. Shahin et al., Hyaluronic acid conjugated nanoparticle delivery of siRNA against TWIST reduces tumor burden and enhances sensitivity to cisplatin in ovarian cancer. Nanomedicine 14, 1381–1394 (2018).

46. T. John, G. Liu, M. S. Tsao, Overview of molecular testing in non-small-cell lung cancer: mutational analysis, gene copy number, protein expression and other biomarkers of EGFR for the prediction of response to tyrosine kinase inhibitors. Oncogene 28 Suppl 1, S14–23 (2009).

47. N. Fedchenko, J. Reifenrath, Different approaches for interpretation and reporting of immunohistochemistry analysis results in the bone tissue - a review. Diagn Pathol 9, 221 (2014).

48. A. Dey et al., Evaluating the Mechanism and Therapeutic Potential of PTC-028, a Novel Inhibitor of BMI-1 Function in Ovarian Cancer. Mol Cancer Ther 17, 39–49 (2018).

49. L. F. Escobar-Hoyos et al., Altered RNA Splicing by Mutant p53 Activates Oncogenic RAS Signaling in Pancreatic Cancer. Cancer Cell 38, 198–211 e198 (2020).

