## Supplementary Figure for "Activation of ERK by altered RNA splicing in cancer"

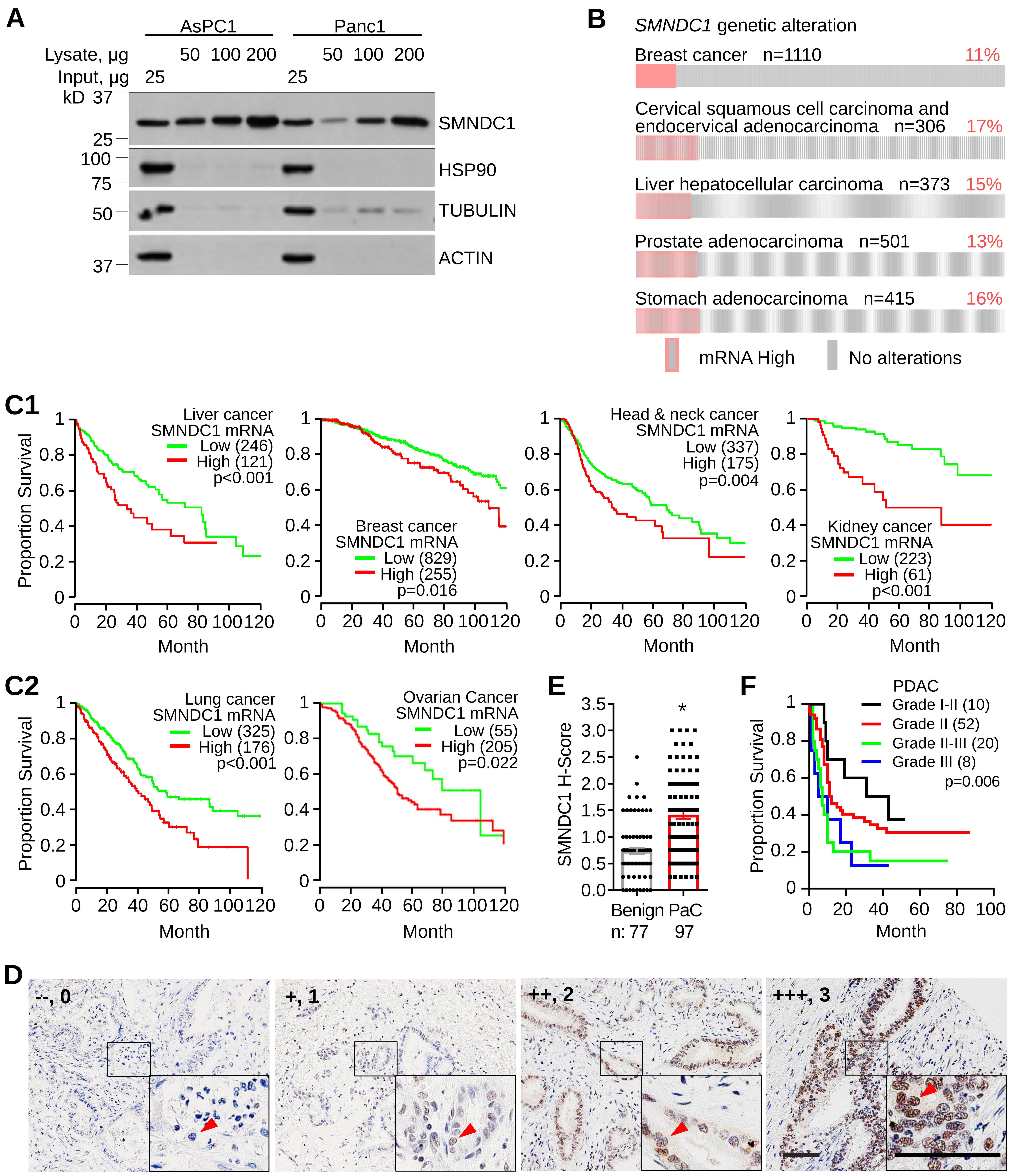


**Supplementary Figure 1.** SMNDC1 is enriched by gold nanoparticles (GNP) and associated with cancer. **A,** Cell lysates from AsPC1 and Panc1 cells were incubated with GNP (40 µg/mL). GNPs binding proteins were boiled off and loaded for WB. 25 µg original lysate proteins were used as Input controls. ACTIN, TUBULIN and HSP90 were used as loading and binding controls. **B.** Oncoprint visualizing SMNDC1 genetic alteration in multiple cancer types. n: case number. **C,** Association of SMNDC1 mRNA expression with the survival of patients with cancer of liver, breast, head and neck, kidney or lung based on TCGA database, and ovary based on another GEO dataset (GSE32062). **D,** Intensity definition of TMA staining. Arrow heads: cancer cell nucleus (red). Scale bar: 100 µm. **E,** TMA H-score for all types of PDAC combined. n, case number. **F,** Association of tumor grade with patient survival based on PDAC TMA clinical data. Number in parentheses: case number. *, p<0.05.

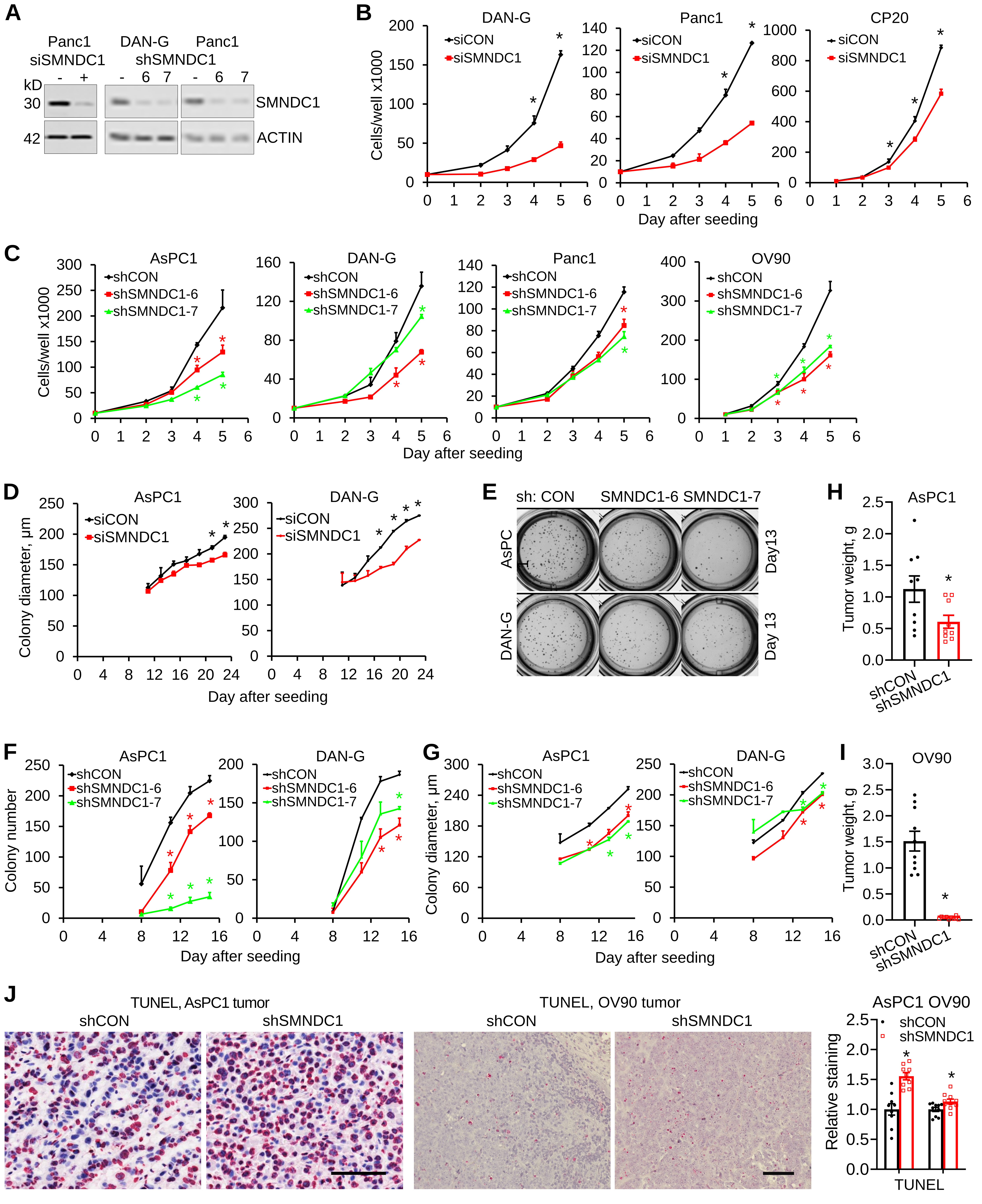


**Supplementary Figure 2.** SMNDC1 promotes cancer cell proliferation, colony formation and tumor growth in mice. **A,** Efficiency of SMNDC1 knockdown by siRNA and shRNA. shRNAs targeting two different sequences of SMNDC1, designated as shSMNDC1-6 and shSMNDC1-7, were used. ACTIN was used as a loading control. **B,** Proliferation of cancer cells upon transient silencing of SMNDC1. **C,** Proliferation of cancer cells with stable silencing of SMNDC1. **D,** 3D colony formation (colony size) of cells upon transient silencing of SMNDC1. **E, F, G,** Example images, colony number and colony size of 3D colony formation assay using stable silencing cells. **H, I,** AsPC1 and OV90 xenograft tumor weight upon experiment termination. **J,** TUNEL staining of AsPC1 and OV90 xenograft tumors and quantification. Scale bar: 100 µm. *, p<0.05.

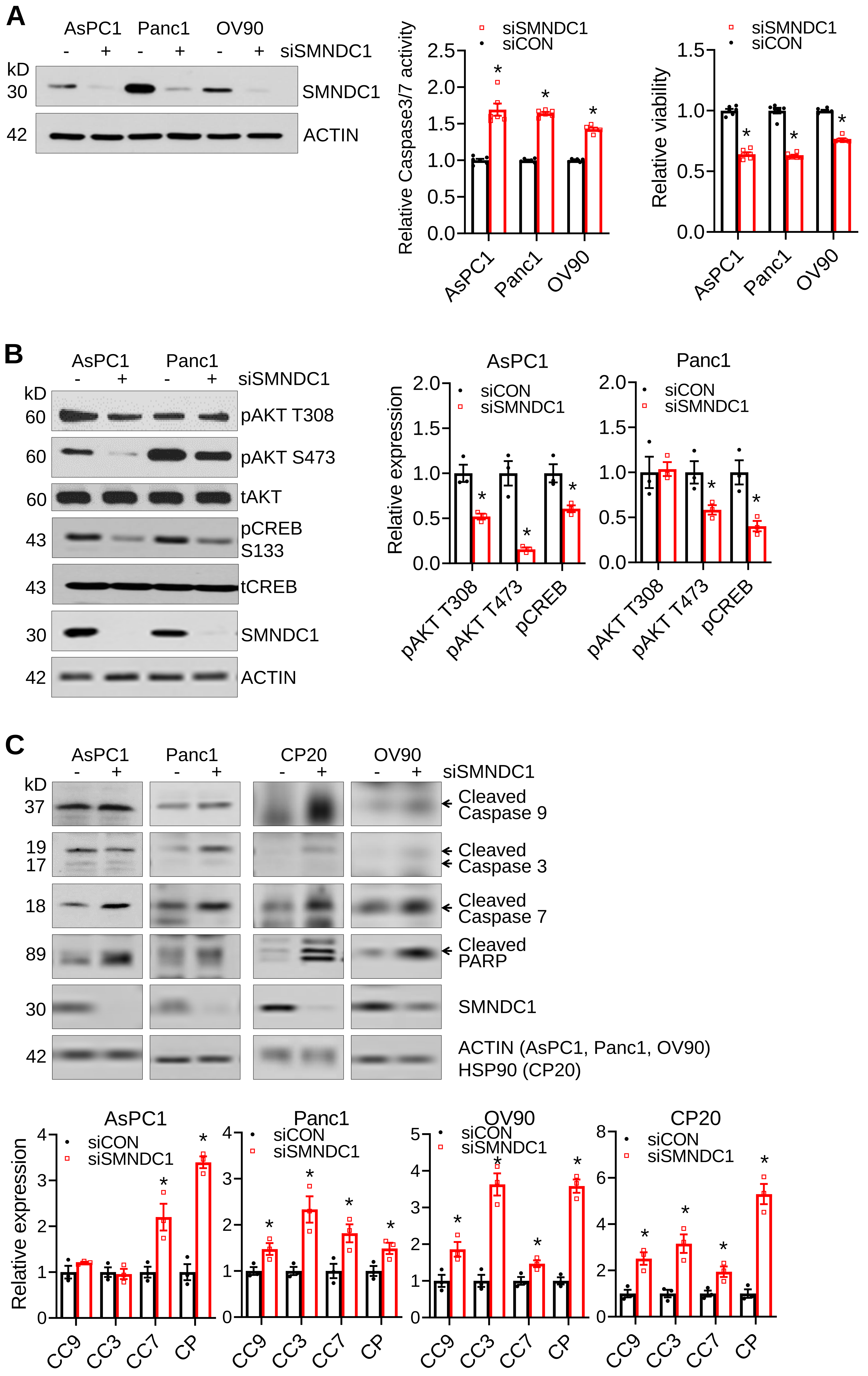


**Supplementary Figure 3.** SMNDC1 upregulates the ERK pathway and blocks pro-apoptotic pathways. **A,** Cell apoptosis, measured by caspase3/7 activity, and survival, measured by glycyl-phenylalanyl-aminofluorocoumarin (GF-AFC) live-cell protease activity, 96 h after siSMNDC1 transfection. Efficiency of SMNDC1 silencing was shown on the left. **B,** Phosphorylation confirmation of chosen kinases based on phospho-kinase array. **C,** Cleavage of caspases and PARP in common intrinsic apoptosis pathway in SMNDC1 knockdown cells and quantification of relative expression. *, $p<0.05$.

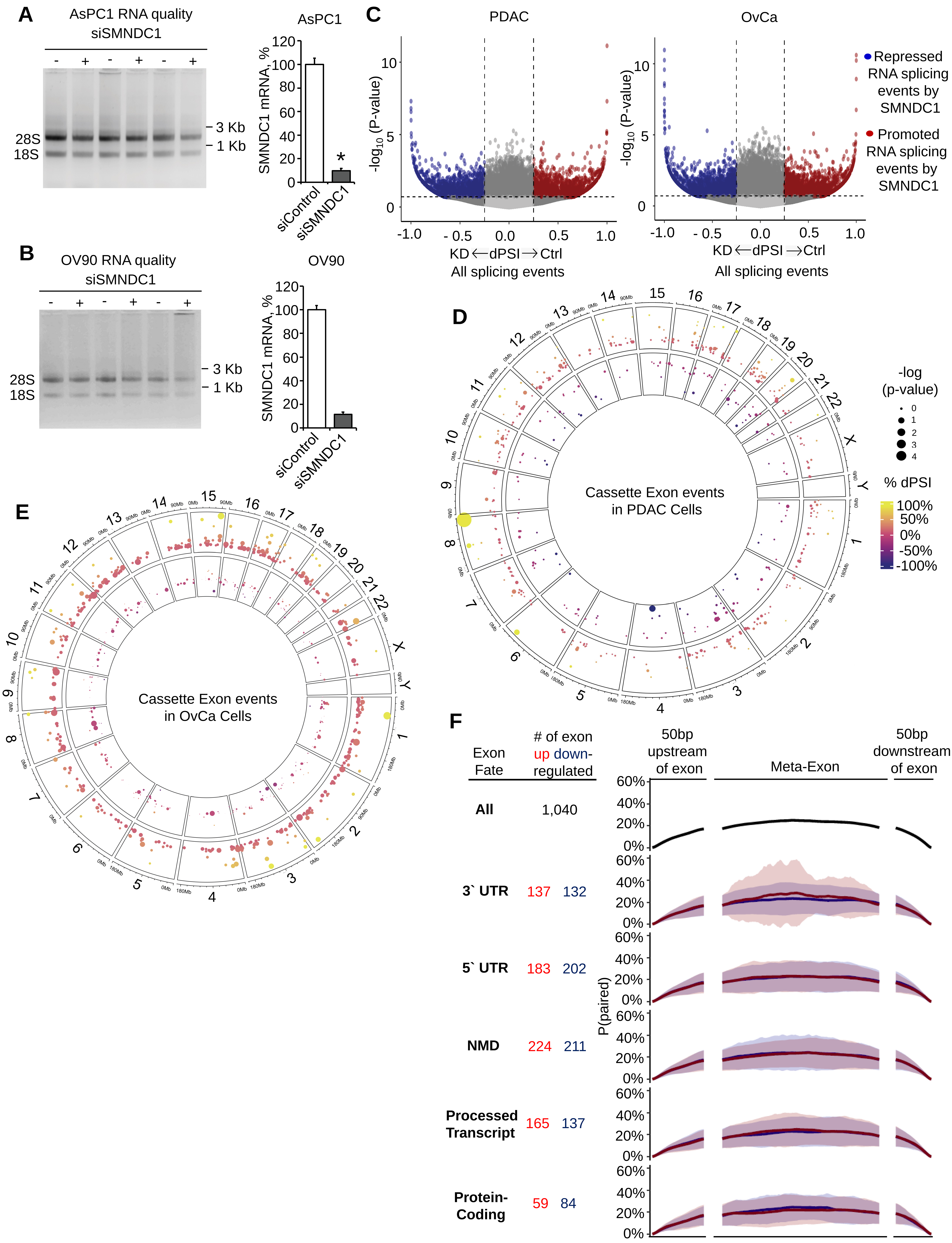
A
AsPC1 RNA quality
siSMNDC1
- + - + - +
28S
18S
3 Kb
1 Kb
AsPC1
SMNDC1 mRNA, %
120
100
80
60
40
20
0
*
siControl
siSMNDC1
B
OV90 RNA quality
siSMNDC1
- + - + - +
28S
18S
3 Kb
1 Kb
OV90
SMNDC1 mRNA, %
siControl
siSMNDC1
C
PDAC
OvCa
-log10 (P-value)
10
5
0
-1.0
- 0.5
0.0
0.5
1.0
KD ← dPSI → Ctrl
All splicing events
Repressed RNA splicing events by SMNDC1
Promoted RNA splicing events by SMNDC1
D
Cassette Exon events in PDAC Cells
-log (p-value)
0
1
2
3
4
% dPSI
100%
50%
0%
-50%
-100%
E
Cassette Exon events in OvCa Cells
F
Exon Fate
### of exon up down-regulated
50bp upstream of exon
Meta-Exon
50bp downstream of exon
P(paired)
60%
40%
20%
0%
All
1,040
3` UTR
137
132
5` UTR
183
202
NMD
224
211
Processed Transcript
165
137
Protein-Coding
59
84

**Supplementary Figure 4.** SMNDC1 promotes inclusion of guanine/cytosine-rich cassette exons, impacting expression of ERK1. **A,** RNA quality for AsPC1 and (**B**) OV90 cells and corresponding SMNDC1 mRNA level after transient knockdown of SMNDC1. **C,** Volcano plot represent the repressed (Blue) and promoted (Red) splicing events in PDAC and OvCa by SMNDC1. Here, KD, SMNDC1 Knockdown cell lines, Ctrl, Control cell lines, dPSI, difference in percentage spliced in. **D,** Circus plots of PDAC and OvCa (**E**) demonstrate the up and down regulated cassette exon (CE) events in respective chromosomes positions. Here, each bubble represents the CE event, whereas the color represents the dPSI, and the size of the bubble represents the significance of the CE event. **F,** Meta-exon plot showing probability of secondary structure formation significantly up/down-regulated CE events categorized by their exon fates: SMNDC1 preferentially recognizes and retains 3'-UTR exons with higher probability of secondary structure formation. p(paired) describes the probability of a given base along the exon sequence to base-pair with another base, predicted by RNAfold at biological temperature (37º C). *, $p<0.05$.

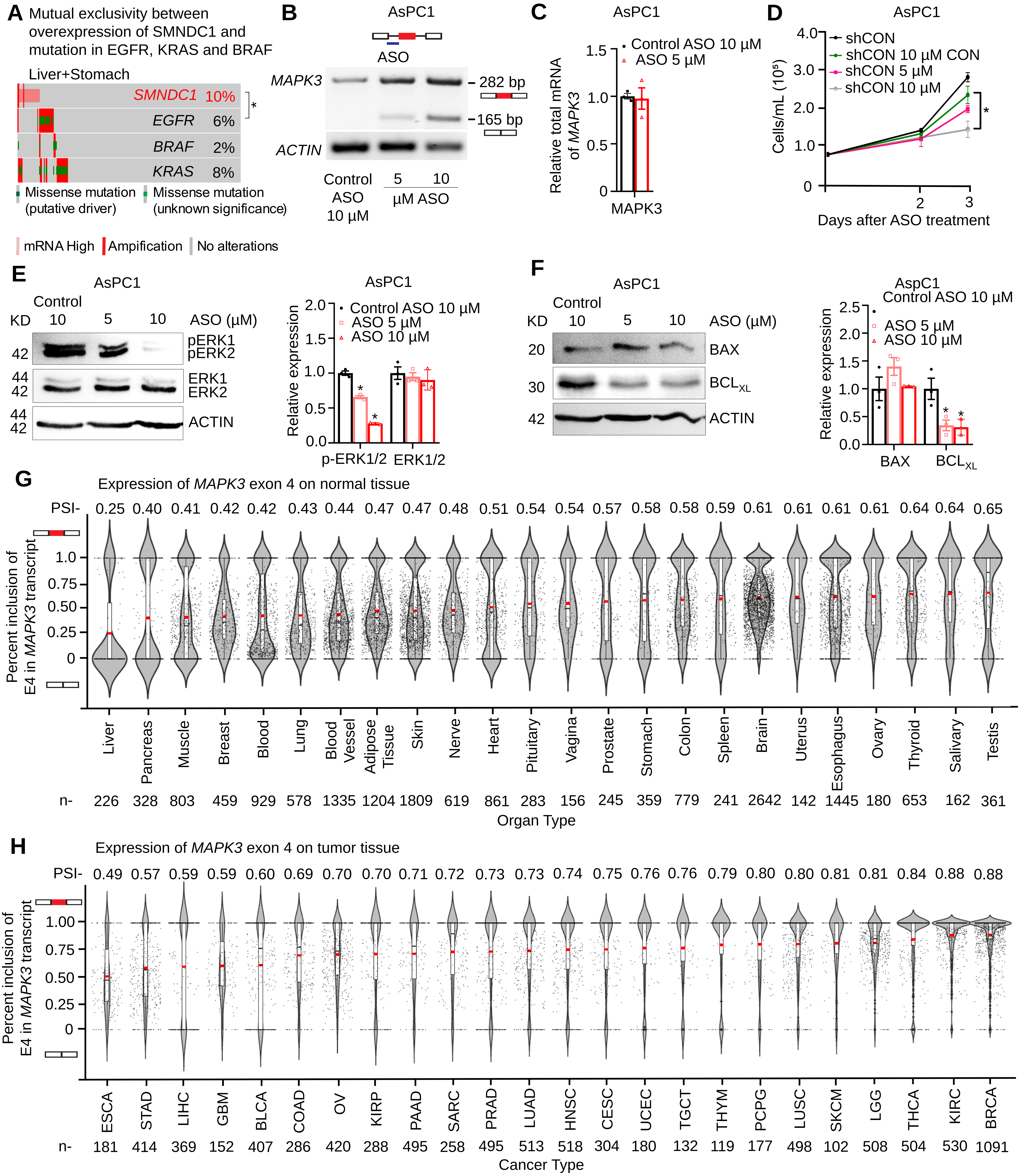


**Supplementary Figure 5.** Inclusion of exon 4 (E4) of MAPK3 (ERK1) by SMNDC1 promotes cell proliferation and increased activity of ERK1 in AsPC1 cells. **A**. Amplification and missense mutations in EGFR/KRAS/BRAF is mutually exclusive to the overexpression of SMNDC1 in liver+stomach cohorts. **B,** Morpholino antisense oligonucleotides (ASOs) block *MAPK3* E4 inclusion in AsPC1 cells. **C,** Total RNA level for *MAPK3* after ASO treatment. **D,** Proliferation curve of ASPc1 cells with and, without ASOs. **E,** Western blots for phospho-ERK1/2, ERK1/2, (**F**) BAX (pro-apoptotic) and $BCL_{XL}$ (anti-apoptotic) upon skipping of E4 in *MAPK3* by ASOs in AsPC1 cells. **G**, Percent spliced in (PSI) value of MAPK3 Exon 4 on normal tissue and (**H**) tumor tissue. *, $p<0.05$
